# Multi-selective RAS(ON) Inhibition Targets Oncogenic RAS Mutations and Overcomes RAS/MAPK-Mediated Resistance to FLT3 and BCL2 Inhibitors in Acute Myeloid Leukemia

**DOI:** 10.1101/2025.06.10.658786

**Authors:** Bogdan Popescu, Matthew F. Jones, Madison Piao, Elaine Tran, Andrew Koh, Isabelle Lomeli, Cheryl A.C. Peretz, Natalia Murad, Sydney Abelson, Carolina Morales, Jose M. Rivera, Yana Pikman, Michael L. Cheng, Aaron C. Logan, Elliot Stieglitz, Catherine C. Smith

**Affiliations:** Department of Medicine, Division of Hematology/Oncology, University of California San Francisco, San Francisco, CA, USA; Department of Pediatrics, Division of Oncology, University of California San Francisco, San Francisco, CA, USA; Department of Pediatric Oncology, Dana-Farber Cancer Institute, Division of Hematology/Oncology, Boston Children’s Hospital and Harvard Medical School, Boston, MA, USA

**Author notes:** **Correspondence to:** Catherine C. Smith, Division of Hematology/Oncology, Department of Medicine, University of California San Francisco, 505 Parnassus Ave, Box 1270 94143, San Francisco, California, USA.

## Abstract

Aberrant activation of the RAS/MAPK signaling limits the clinical efficacy of several targeted therapies in acute myeloid leukemia (AML). In *FLT3*-mutant AML, the selection of clones harboring heterogeneous RAS mutations drives resistance to FLT3 inhibitors (FLT3i). RAS activation is also associated with resistance to other AML targeted therapies, including the BCL2 inhibitor venetoclax. Despite the critical need to inhibit RAS/MAPK signaling in AML, no targeted therapies have demonstrated clinical benefit in RAS-driven AML. To address this unmet need, we investigated the preclinical activity of RMC-7977, a multi-selective inhibitor of GTP-bound active [RAS(ON)] isoforms of mutant and wild-type RAS in AML models. RMC-7977 exhibited potent antiproliferative and pro-apoptotic activity across AML cell lines with MAPK-activating signaling mutations. In cell line models with acquired FLT3i resistance due to secondary RAS mutations, treatment with RMC-7977 restored sensitivity to FLT3i. Similarly, RMC-7977 effectively reversed resistance to venetoclax in RAS-addicted cell line models with both RAS wild-type and mutant genetic backgrounds. In murine patient-derived xenograft models of RAS-mutant AML, RMC-7977 was well tolerated and significantly suppressed leukemic burden in combination with gilteritinib or venetoclax. Our findings strongly support clinical investigation of broad-spectrum RAS(ON) inhibition in AML to treat and potentially prevent drug resistance due to activated RAS signaling.

## Introduction

Acute myeloid leukemia (AML) is an aggressive hematologic malignancy characterized by clonal expansion and abnormal differentiation of myeloid progenitor cells. Despite decades of research, the prognosis of patients with relapsed or treatment-refractory (R/R) disease remains very poor.^1^ Therapies targeted at molecular vulnerabilities of AML, such as inhibitors of Fms-Like Tyrosine Kinase-3 (FLT3), B-cell lymphoma 2 (BCL2), or isocitrate dehydrogenase (IDH) 1/2, have improved outcomes for defined subgroups of patients.^2–9^ However, remissions achieved with these therapies are often transient, mainly due to their vulnerability to primary and adaptive resistance. Mutations in Rat Sarcoma Virus (RAS) family oncogenes (predominantly in *NRAS* and *KRAS*) are observed in approximately 10-30% of AML cases.^10,11^ Although the prognostic significance of RAS mutations has varied across studies, and only *KRAS* mutations have been associated with adverse outcomes^12–15^, aberrant activation of the RAS/MAPK signaling pathway is commonly involved in resistance to multiple AML targeted therapies.

In AML with *FLT3* mutations (*FLT3*-mut), FLT3 tyrosine kinase inhibitors (FLT3i), such as midostaurin, quizartinib, and gilteritinib, are now standard of care for both newly diagnosed and R/R disease, but patients treated with single-agent FLT3i invariably relapse despite initial clinical response.^16^ Multiple, heterogeneous clones harboring mutations that bypass FLT3 inhibition and reactivate downstream signaling of the RAS/MAPK pathway (*NRAS*, *KRAS*, *PTPN11*) have been identified at relapse in patients who received FLT3i.^17–20^ We have previously shown using serial DNA single-cell sequencing studies that RAS mutations are often present at a subclonal level prior to treatment and expand under the selective pressure of FLT3i, as both *FLT3* wild-type and *FLT3*/RAS co-mutant resistant clones.^18,21^

The BCL2 inhibitor (BCL2i) venetoclax in combination with hypomethylating agents (HMA) has become standard of care for newly diagnosed unfit patients with AML.^9^ In FLT3-mut AML, the combination of gilteritinib and the BCL2 inhibitor (BCL2i) venetoclax has superior response rates compared to gilteritinib monotherapy.^8^ Nonetheless, primary and acquired resistance to BCL2i combinations remain common. Retrospective studies of patients receiving HMA and venetoclax have linked venetoclax resistance to RAS mutations or to AML with monocytic features.^22–25^ Interestingly, RAS mutations and RAS signaling activation have also been correlated with monocytic differentiation.^26–29^ Putative mechanisms for RAS-mediated venetoclax resistance include increased expression and stability of MCL1, causing decreased dependence on BCL2, mitochondrial membrane remodeling, and metabolic reprogramming with increased levels of oxidative phosphorylation.^22,30–32^ A venetoclax-resistant monocytic differentiation state with higher MCL1 and lower BCL2 expression has been found to arise from a subpopulation of RAS-mutant leukemia stem cells (LSCs), and monocytic differentiation is generally associated with venetoclax resistance.^29,33^ However, our group has recently identified that RAS signaling activation can be upregulated during treatment with gilteritinib and venetoclax to drive monocytic differentiation and resistance even in RAS wild-type clones.^21^

Given the poor prognosis of R/R AML, the dearth of effective treatments for these patients, and the central role of RAS mutations in driving resistance to targeted therapies, new approaches to target RAS signaling are urgently needed. Historically, RAS was deemed “undruggable” due to its protein structure lacking deep, hydrophobic pockets amenable to small-molecule binding. ^34^ Early efforts to disrupt downstream signaling with MEK inhibitors (MEKi) or to interfere with RAS farnesylation have not yielded clinically useful treatments in AML.^35–39^ More recently, direct inhibitors of specific *KRAS*-mutant oncoproteins have shown promise in solid tumors, demonstrating that direct RAS inhibition is feasible.^40–45^ Still, the heterogeneity of RAS-mutant clones identified in AML limits the utility of isoform- or mutation-specific inhibitors and highlights the need for broad-spectrum RAS inhibition.

RMC-7977 is a potent, orally bioavailable small molecule inhibitor of both wild-type and mutant RAS proteins in the GTP-bound active state [RAS(ON) multi-selective]^46^ and is a preclinical tool compound representative of the investigational drug daraxonrasib (RMC-6236), currently being tested in a set of clinical trials.^47^ RMC-7977 non-covalently binds to the intracellular chaperone cyclophilin A, generating a neomorphic interface with high affinity for all RAS isoforms. The resulting tri-complexes sterically block RAS-effector interactions required for propagating oncogenic signals.^46^ In Phase 1/1b trials, daraxonrasib showed encouraging clinical activity and an acceptable safety profile in patients with advanced solid tumors. Phase 3 trials of daraxonrasib, RASolute 302 (NCT06625320) and RASolve 301 (NCT06881784) as second-line therapy in metastatic pancreatic ductal adenocarcinoma (PDAC) and non-small cell lung cancer (NSCLC), respectively, are currently ongoing.^48–51^

In this study, we investigate the preclinical activity of the RAS(ON) multi-selective inhibitor RMC-7977 in AML models with aberrant RAS/MAPK signaling, including clinically relevant models of AML resistant to FLT3 and/or BCL2 inhibitors. RMC-7977 demonstrates potent single-agent activity in AML cell lines driven by constitutively active RAS mutations and mutations in upstream receptor tyrosine kinases (RTKs). RMC-7977 also re-sensitizes *FLT3* and RAS co-mutated AML to the antileukemic activity of gilteritinib and inhibits the outgrowth of gilteritinib-resistant RAS-mutant clones. Furthermore, we show that RAS(ON) inhibition reverses resistance in venetoclax-resistant AML characterized by increased monocytic differentiation and RAS signaling activation. Our findings provide a strong rationale for the clinical investigation of RAS(ON) inhibitors in combination with established targeted therapies to improve outcomes in patients with RAS-driven AML.

## Results

### RMC-7977 is active in cell lines driven by RAS/MAPK-activating mutations

Because the RAS(ON) multi-selective inhibitor RMC-7977 showed potent activity in solid tumor cell lines driven by activating RAS mutations^46,52^, we hypothesized that RMC-7977 would be active in AML cell lines highly dependent on RAS/MAPK signaling. To test this hypothesis, we treated seven AML cell lines with a range of RMC-7977 concentrations and measured cell viability. These cell lines harbor activating mutations in *NRAS*^Q61L^ (OCIAML-3, HL-60), and *KRAS*^G13D^ (NOMO-1), and in upstream receptor tyrosine kinases (RTK) known to activate the MAPK pathway: *FLT3*-ITD (Molm-14, MV4-11) and *KIT*^N822K^ (Kasumi-1, SKNO-1). RMC-7977 potently inhibited cell proliferation in all cell lines at IC_50_ concentrations in the low nanomolar range (Figure 1A-C). In contrast, the HEL cell line, driven by the *JAK2*^V612F^ mutation, was insensitive to RMC-7977 (Supplementary figure S1A), confirming that the antiproliferative activity is specific to cell lines dependent on MAPK signaling for proliferation. To understand the biochemical effects of RMC-7977 treatment on RAS signaling, we assayed downstream protein phosphorylation by western blotting and observed potent, concentration-dependent inhibition of ERK phosphorylation in all cell lines tested (Figure 1D). We also noted a partial, cell line-dependent inhibition of AKT phosphorylation, suggesting RAS-mediated inhibition of the PI3K pathway. This finding is in agreement with the previous observation that RMC-7977 inhibits PI3Kα binding to RAS proteins.^46^ Consistent with ERK and AKT signaling downregulation, RMC-7977 inhibited phosphorylation of ribosomal protein S6 at serine residues 235/236 in each cell line, albeit to varying degrees (Figure 1D).

**Figure 1.**
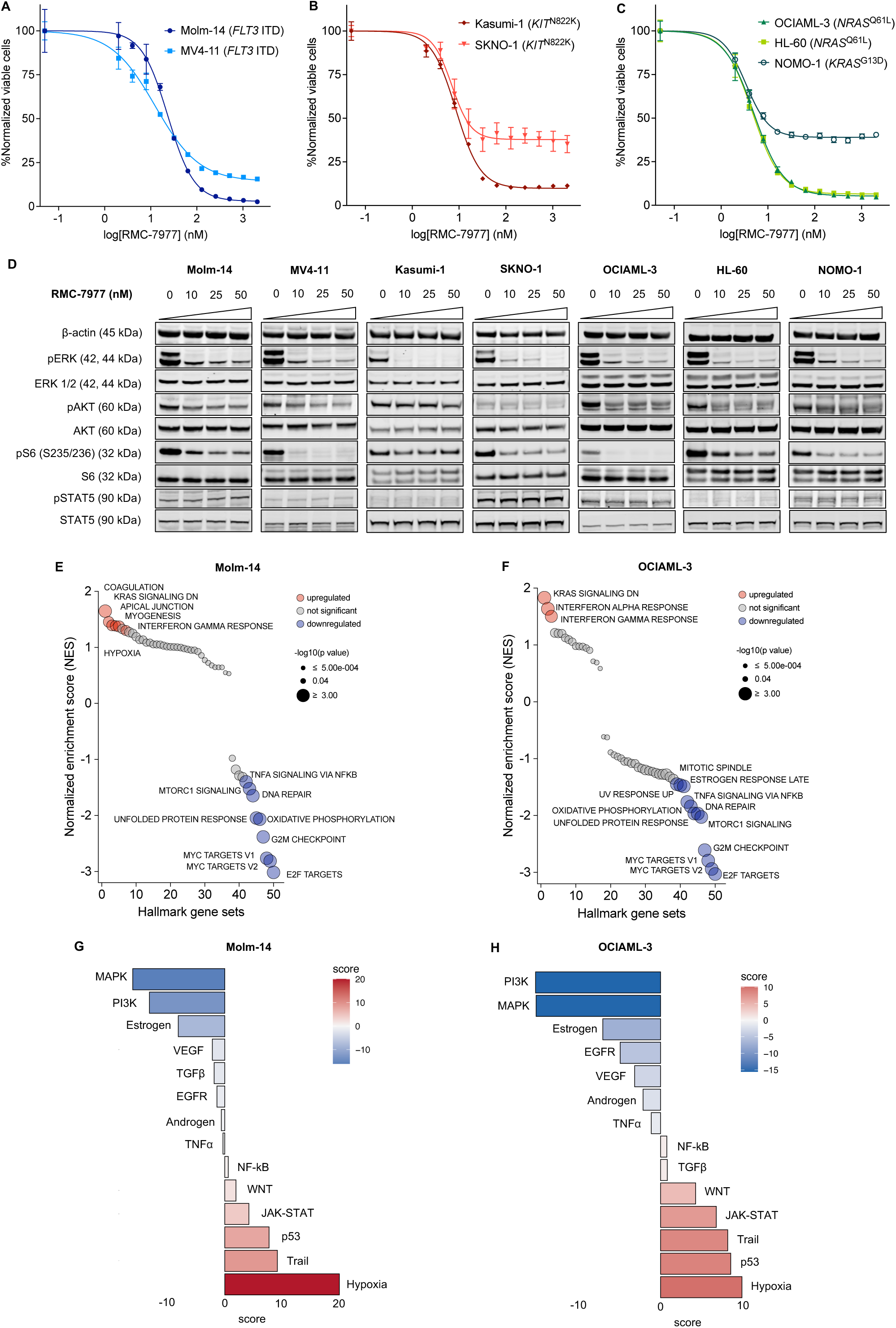
Activity of RAS(ON) multi-selective inhibitor RMC-7977 in AML cell lines. A-C: Concentration-response curves representing relative proliferation of AML cell lines driven by *FLT3* (A), *KIT* (B), *NRAS* or *KRAS* mutations (C) after 48 h of exposure to serial doses of RMC-7977. Data represent the mean ± SD of three replicates. D: Western blot analysis of seven AML cell lines exposed for 2 hours to indicated concentrations of RMC-7977; β-actin was used as loading control. E-F: Bubble plots showing enrichment of GSEA Hallmark gene sets expression in Molm-14 and OCIAML-3 cell lines after exposure for 24 hours to RMC-7977 (25 nM). The size of the bubbles represents statistical significance, and the colors indicate upregulated (red) or downregulated (blue) expression. G-H: PROGENy pathway activity scores in Molm-14 and OCIAML-3 cells after exposure for for 24 hours to RMC-7977 (25 nM).

To investigate the transcriptional changes that occur with RAS(ON) inhibition, we performed bulk RNA sequencing (RNA-seq) in two RAS/MAPK-activated cell lines driven by either *FLT3*-ITD (Molm-14) or *NRAS*^Q61L^ mutation (OCIAML-3) after treatment with RMC-7977 at an early (2 hours) and a late time point (24 hours). In both cell lines, RMC-7977 altered the expression of a higher number of genes (|log_2_FC| ≥ 2, p-value < 0.05) at 24 hours than at 2 hours (Supplementary Figure S1B-C), suggesting a time-dependent transcriptional response. Gene Set Enrichment Analysis (GSEA) of Hallmark gene sets (MSigDB)^53^ showed a rapid decrease in the expression of genes upregulated by *KRAS* signaling, downregulation of Myc targets, as well as downregulation of several inflammatory gene sets (Inflammatory response, IL6/STAT3, TNFα via NF-kB) (Supplementary Figure S1D-E). At 24 hours, in addition to a persistent downregulation of RAS transcriptional targets, we also observed simultaneous downregulation of genes involved in mTORC1 signaling, cell cycle and DNA repair, mitotic spindle, protein cellular stress response or oxidative phosphorylation, as well as prominent upregulation of interferon-responsive genes (Figure 1E-F). To infer signaling pathway activity from RNAseq data, we performed Pathway RespOnsive GENes (PROGENy)^54^ analysis. In both cell lines treated with RMC-7977 we noted a marked and persistent decrease in activity of both MAPK and PI3K pathways. Additionally, treatment with RMC-7977 led to enrichment of pathway activity scores corresponding to hypoxia, p53-mediated DNA damage response, and Trail pathway of apoptosis, with a more pronounced effect at a later time point. RMC-7977 treatment also led to a decreased activity score of estrogen pathway (Figure 1G-H, Supplementary Figure S1F-G). Although the roles of estrogen signaling in myeloid malignancies are not fully elucidated, reports have demonstrated *in vitro* pro-apoptotic activity of selective estrogen receptor modulators (SERM) in AML cells.^55^ In keeping with previous reports in solid malignancies^56^, we also observed increased JAK/STAT signaling activity, a potential adaptive response to RAS inhibition. In summary, we found that RMC-7977 induces a series of anti-proliferative and pro-apoptotic transcriptional responses consistent with broad RAS effector pathway inhibition.

### RMC-7977 induces apoptosis in AML cell lines via dual inhibition of RAS/MAPK and PI3K/AKT signaling pathways

To assess the ability of RMC-7977 to induce apoptotic cell death in AML cell lines, we performed a flow cytometry-based caspase activation assay in RTK and RAS-driven cell lines evaluated in Figure 1A-C. In contrast to anti-proliferative effects, there was more variability in the potency of pro-apoptotic activity across cell lines (Figure 2A). To compare the pro-apoptotic effects of RAS/MAPK pathway inhibition at different levels, we performed a comparative caspase assay with the RAS(ON) inhibitor and several downstream MAPK pathway inhibitors - the RAF inhibitor naporafenib, the MEK inhibitor trametinib, and the ERK inhibitor ulixertinib. In the cell lines highly sensitive to apoptosis Molm-14 and OCIAML-3, RMC-7977 induced higher levels of apoptosis at lower doses compared to downstream inhibitors (Figure 2B-C). Although the activated caspase 3/7 levels observed with RMC-7977 treatment were less significant in the rest of the AML cell lines tested, they were generally higher than those observed with the downstream inhibitors in most cell lines, except for the *KIT*-mutant cell line SKNO-1 (Supplementary figure S2A-E). These findings suggest that upstream RAS/MAPK pathway inhibition elicits a higher apoptotic response compared to downstream inhibition.

**Figure 2.**
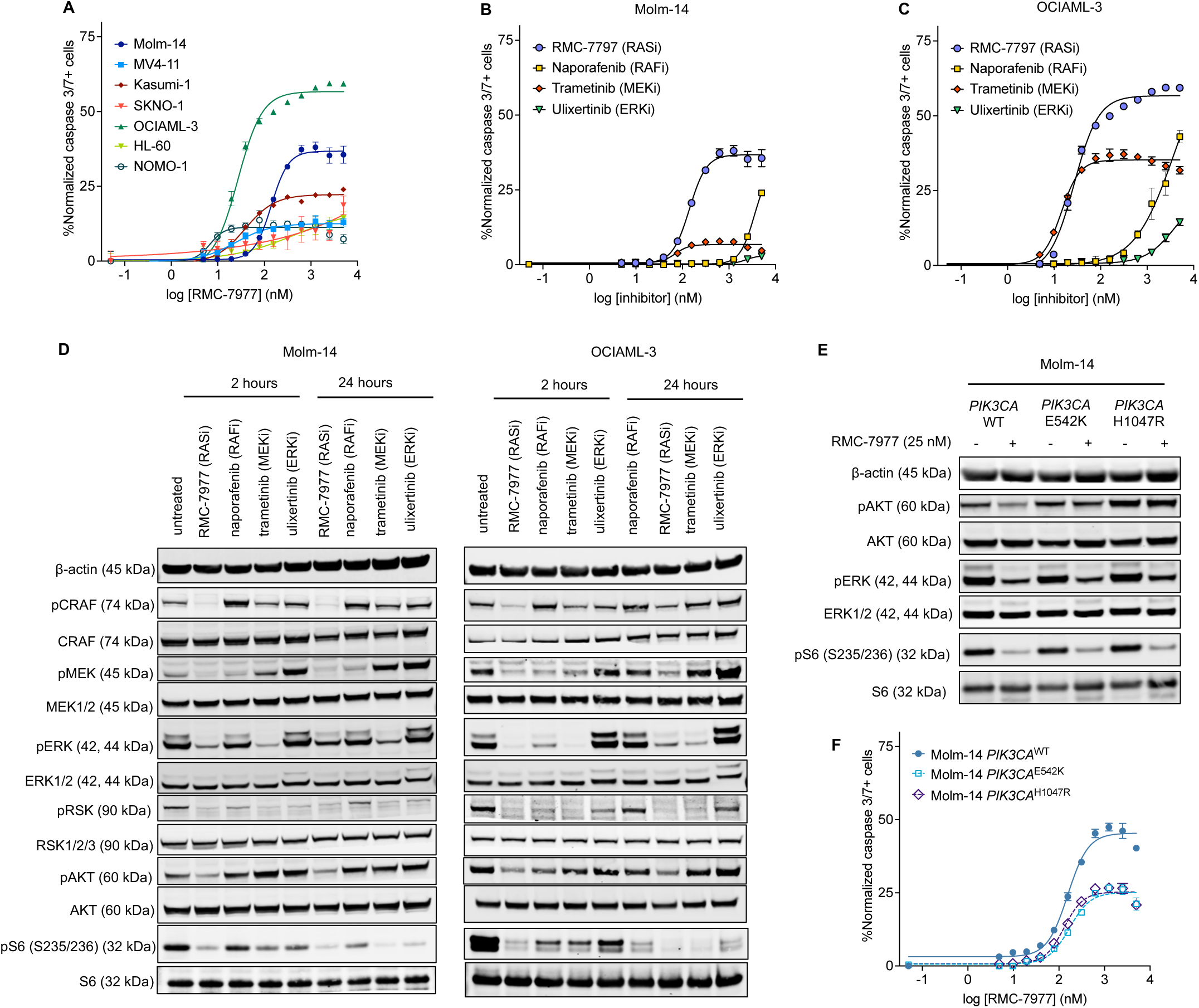
RAS(ON) inhibition has a higher pro-apoptotic activity than downstream MAPK inhibition in AML cell lines. A: Concentration-response curves showing activated caspase 3/7 expression in AML cell lines after 24 h of exposure to increasing doses of RMC-7977, normalized to untreated controls. Data represent the means ± SD of three replicates. B-C: Comparative caspase 3/7 activity in Molm-14 (B) and OCIAML-3 (C) cells exposed to RMC-7977 (RASi), naporafenib (RAFi), trametinib (MEKi) and ulixertinib (ERKi). D-E: Western blot analysis of Molm-14 (D) and OCIAML-3 (E) cells treated for 2 hours and 24 hours with the indicated inhibitors at concentrations approximately equivalent to IC50 for each compound; β-actin was used as loading control. F: Comparative caspase 3/7 activity in Molm-14 cells expressing either *PIK3CA*^WT^ or *PIK3CA*^E542K^/*PIK3CA*^H1047R^ after 24 hours of exposure to RMC-7977. Data represent the means ± SD of three replicates.

To characterize changes in MAPK signaling after inhibition of upstream versus downstream, we evaluated the phosphorylation of MAPK substrates at different levels of the pathway after early (2 hours) and late (24 hours) exposure to treatment with RMC-7977, naporafenib, trametinib, and ulixertinib at IC_50_ concentrations for each compound. As expected, in Molm-14 and OCIAML-3 cells, all compounds persistently reduced the phosphorylation of their downstream targets (Figure 2D). We observed a rebound effect in MEK, ERK, and RSK phosphorylation at 24 hours relative to the 2-hour timepoint with naporafenib, but only a minor rebound with the other inhibitors. Compared to other treatments, RMC-7977 induced partial inhibition of AKT phosphorylation, likely reflective of an impaired interaction between RAS and the PI3K p110α catalytic subunit due to RAS(ON) inhibition, as previously described.^46^ To ascertain whether this RAS-mediated partial inhibition of the PI3K pathway could explain the higher pro-apoptotic effect of RMC-7977 versus downstream inhibitors, we generated Molm-14 cells with doxycycline-inducible overexpression of either *PIK3CA*^WT^ or of two gain-of-function mutations in the helical or kinase domains of PIK3CA, commonly identified in solid malignancies: *PIK3CA*^E542K^ and *PIK3CA*^H1047R^ respectively. Although RMC-7977 downregulated AKT phosphorylation in *PIK3CA*^WT^ cells, it did not in either mutant cell line (Figure 2E). Accordingly, both *PIK3CA*^H1047R^ and *PIK3CA*^E542K^ exhibited a diminished apoptotic response to RMC-7977, compared to *PIK3CA*^WT^ (Figure 2F). We observed a similar reduction in caspase activation in both *PIK3CA* mutants, despite previous data suggesting a lower dependence of the *PIK3CA*^H1047R^ variant on RAS-GTP binding, compared to that of *PIK3CA*^E542K^.^57^ Additionally, to test whether isolated inhibition of either the Raf/MEK/ERK or the PI3K/AKT pathway is sufficient to induce apoptosis, we performed a caspase assay in Molm-14 cells overexpressing a doxycycline-inducible dominant-active MEK allele (*MAP2K1*^S218D/S222D^, referred to as MEK^DD^) or dominant-active AKT variant (*AKT1*^E17K^, Supplementary figure S2F). Similarly to *PIK3CA*-mutant cells, in cells expressing *AKT1*^E17K^, the apoptosis induced by RMC-7977 was attenuated. However, overexpression of MEK^DD^ completely abrogated the apoptotic response (Supplementary figure S2G-H). Collectively, these data indicate that while PI3K/AKT pathway inhibition contributes to apoptosis induced by RMC-7977 in AML cell lines, RAS/MAPK signaling suppression is essential to this effect.

### RMC-7977 overcomes both on- and off-target resistance to FLT3 inhibitors

Given the diversity of RAS mutations that confer clinical resistance to FLT3i, we asked whether broad-spectrum, multi-selective RAS inhibition could address FLT3i resistance driven by selection for heterogeneous RAS-mutant clones. We performed cell viability assays in *FLT3*-ITD^+^ Molm-14 cells that acquired secondary *NRAS*^G12C^ or *NRAS*^Q61K^ mutations during long-term exposure to FLT3i, and we demonstrated that RAS-mutant cells exhibit a variable degree of resistance to gilteritinib compared to the parental cells (Figure 3A). However, co-treatment with RMC-7977 restored gilteritinib sensitivity in both *NRAS*^G12C^ and *NRAS*^Q61K^ mutant cell lines (Figure 3B). As expected, single-agent gilteritinib failed to completely inhibit ERK, AKT and S6 phosphorylation in RAS-mutant cells, whereas the addition of RMC-7977 induced a profound suppression of both phospho-ERK and phospho-AKT (Figure 3C).

**Figure 3.**
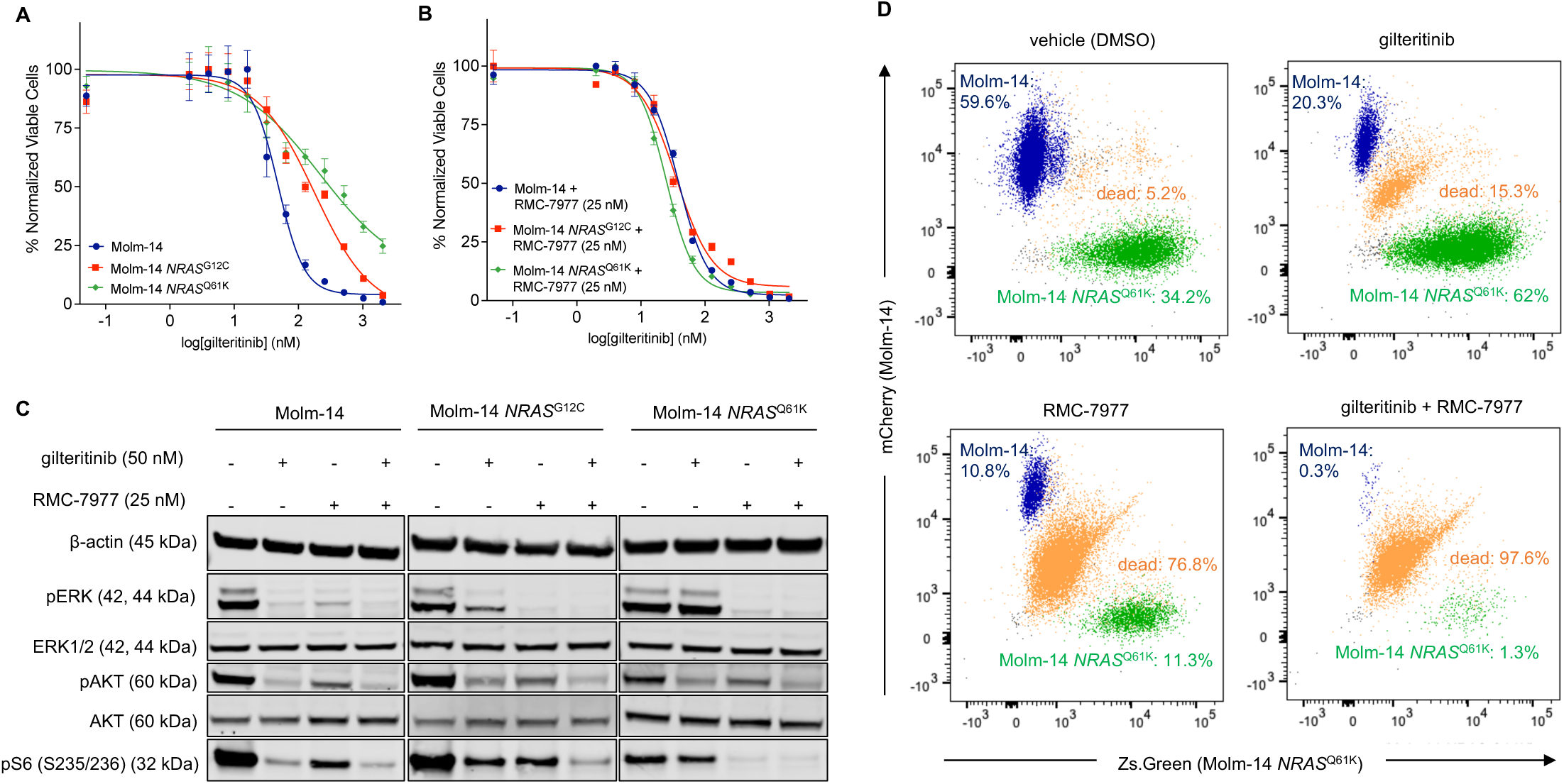
RMC-7977 re-sensitizes *FLT3* and RAS co-mutant AML cells to FLT3 inhibitors. A-B: Concentration-response curves representing relative proliferation of Molm-14 parental cells or Molm-14 expressing secondary *NRAS*^G12C^ or *NRAS*^Q61K^ mutations after 48 h of exposure to serial doses of gilteritinib relative to un-treated controls. In (B), the proliferation assay was conducted in the presence of RMC-7977 (25 nM). Data represent the mean ± SD of three replicates. C: Western blot analysis of parental or *NRAS*-mutant Molm-14 cells treated with either gilteritinib, RMC-7977 or both inhibitors; β-actin was used as loading control. D: Fluorescently-tagged Molm-14 (blue) or Molm-14 *NRAS*^Q61K^ (green) cells were mixed at a 1:1 ratio, then treated with gilteritinib, RMC-7977 or their combination. Scatter plots showing percentages of each cell population after 96 hours of indicated treatments and a staining with a live/dead dye (Sytox AADvanced). Plots representative of three replicates.

Clinical resistance to FLT3i is often caused by the outgrowth of pre-existing subclonal RAS mutations under the selective pressure of FLT3i.^17,18,21^ To model inter-clonal competition and selection *in vitro*, we mixed Molm-14 and Molm-14 *NRAS*^Q61K^ cells tagged with fluorescent proteins at a 1:1 ratio, treated the mixtures with DMSO, gilteritinib, RMC-7977 and their combination for 96 hours, then stained with a live/dead stain (Sytox AADvaced) and assessed viability via flow cytometry (Supplementary figure S3A-B). We observed that in the absence of treatment, the parental cells have a slight growth advantage relative to the *NRAS*^Q61K^ cells, but treatment with gilteritinib inhibited the proliferation of parental cells, allowing for the outgrowth of the RAS co-mutant cell line. RMC-7977 was equally active against both cell populations, while the combination treatment almost entirely suppressed the growth of all leukemia cells in the co-cultured mixed population (Figure 3D).

In addition to the off-target mechanism of resistance represented by RAS mutations, type II FLT3 inhibitors that bind the inactive form of the RTK, such as sorafenib or quizartinib, are also vulnerable to on-target resistance mutations in the tyrosine kinase domain (TKD), predominantly at codons D835 and F691.^58^ We demonstrated that, in contrast to quizartinib, RMC-7977 maintains activity in Molm-14 cells that express secondary *FLT3* TKD mutations (Supplementary figure S3C-D). Taken together, these findings suggest that RMC-7977 effectively targets both on-target and off-target mechanisms of resistance to FLT3i.

### RMC-7977 targets RAS/MAPK-mediated mechanisms of resistance to BCL2 inhibitor venetoclax

Resistance to the BCL2i venetoclax in AML has been linked to both RAS signaling activation and monocytic differentiation, though the causal relationship between RAS genotype and monocytic phenotype remains unclear.^22–25,30^ Recent data suggest that monocytic differentiation in venetoclax-resistant AML is driven by either RAS mutations or RAS/MAPK pathway activation independently of RAS mutational status.^21,33^ To characterize the relationship between monocytic AML, RAS activation, and venetoclax resistance, we interrogated publicly available data from the BEAT AML 2.0 master trial^29^ and identified 335 patient samples with available bulk DNA sequencing, bulk RNAseq, and *ex vivo* drug sensitivity data. We then performed single-sample gene set enrichment analysis (ssGSEA) of the RNAseq data and observed a positive correlation between *ex vivo* resistance to venetoclax (represented by increased AUC values) and enrichment scores of both Hallmark *KRAS* signaling (mSigdb) and monocyte-like^59^ gene sets (Figure 4A, Supplementary figure S4A-C). We also observed correlations between *KRAS* signaling enrichment score and other previously described predictors of response to venetoclax, such as the *BCL2*/*MCL1* expression ratio or expression of *BCL2A1* (Bfl-1) or *BCL2L1* (BCLxL, Figure 4B).^25,60,61^ This analysis isolates a subset of samples with low pharmacologic sensitivity to venetoclax that have a high RAS transcriptional signature, a low *BCL2*/*MCL1* ratio, high *BCL2A1* expression, and transcriptional monocytic features. Although RAS mutations themselves are associated with resistance to BCL2i^25^ (Supplementary figure S4D), we also observed many AML samples that lack any RAS mutation yet shared similar drug sensitivity and transcriptional profiles. After excluding the samples with identifiable RAS mutations by bulk DNA sequencing, we also observed similar correlations in the RAS^WT^ subset of samples (Supplementary figure S4E-F). This finding suggests that the RAS/MAPK transcriptional activation can drive the monocytic cell state associated with resistance to BCL2i independent of RAS mutational status.

**Figure 4.**
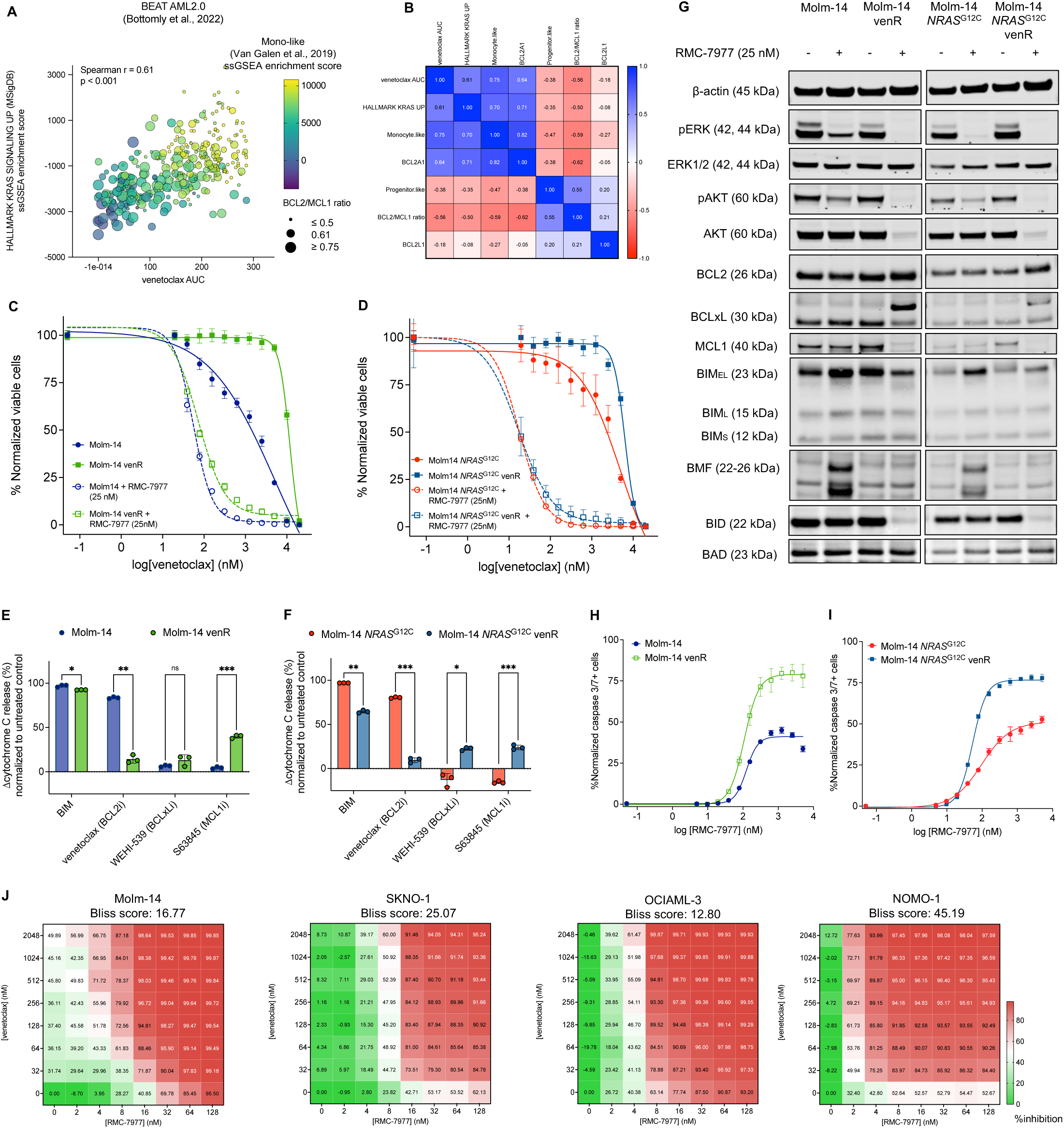
RMC-7977 inhibits RAS activation driving venetoclax resistance in both RAS^mut^ and RAS^WT^ AML. A: Bubble plot showing correlation between *ex vivo* sensitivity to venetoclax (AUC) and RAS transcriptional signature (Hallmark *KRAS* Signaling Up^53^, ssGSEA enrichment score) in 335 RAS^WT^ and RAS^mut^ primary AML samples from the BEAT AML2.0 trial.^29^ Size of bubbles represent *BCL2*/*MCL1* normalized expression ratio and color gradient represents transcriptional scores of monocytic differentiation (Monocyte-like^59^, ssGSEA enrichment score). B: Correlation matrix of nonparametric Spearman r correlation coefficients between venetoclax *ex vivo* sensitivity and indicated transcriptional parameters. C-D: Comparative concentration-response curves representing relative proliferation of Molm-14 and venetoclax-resistant Molm-14 cells (C) or Molm-14 *NRAS*^G12C^ and venetoclax-resistant Molm-14 *NRAS*^G12C^ cells (D) after 48 h of exposure to serial doses of venetoclax with or without RMC-7977 (25 nM). E-F: Dynamic iBH3 profiling showing relative differences in cytochrome c release upon exposure to BH3-mimetic peptide (BIM) or inhibitors (venetoclax, WEHI-539 and S63845) in Molm-14 (E) or Molm-14 *NRAS*^G12C^ (F) cells, compared to their venetoclax-resistant counterparts. Data represent the mean ± SD of three replicates; unpaired Welch t-test with Holm-Sidak’s correction for multiple comparisons was used for statistical significance (∗∗∗∗p ≤ 0.0001, ∗∗∗p ≤ 0.001, ∗∗p ≤ 0.01, ∗p ≤ 0.05, ns, non-significant). H-I: Comparative activated caspase 3/7 expression between venetoclax-sensitive and resistant Molm-14 (H) or Molm-14 *NRAS*^G12C^ cells after 24 hours of exposure to RMC-7977. Data represent the means ± SD of three replicates. J: Concentration-response matrices representing normalized cell viability inhibition following 48 h of treatment with increasing doses of RMC-7977 and venetoclax in the indicated cell lines. Synergy scores were computed using the Bliss method within SynergyFinder v.3.0 software.

We reasoned that if the venetoclax-resistant monocytic phenotype is driven by hyperactive RAS signaling, then RMC-7977 may offer a tool to restore venetoclax sensitivity in both RAS^mut^ and RAS^WT^ venetoclax-resistant AML cells. First, using cell viability assays, we established that Molm-14 cells with doxycycline-inducible overexpression of *NRAS*^G12C^ and *NRAS*^Q61K^ mutations have a lower sensitivity to venetoclax relative to Molm-14 cells overexpressing *NRAS*^WT^, but co-treatment with RMC-7977 re-sensitized both RAS^mut^ cell lines to BCL2 inhibition (Supplementary figure S4G-H).

Next, we generated venetoclax-resistant (venR) cell lines with both RAS^WT^ (Molm-14) and RAS^mut^ (Molm-14 *NRAS*^G12C^) genetic backgrounds after long-term exposure to incrementally increasing concentrations of venetoclax. No acquired mutations in RAS or other genes along the RAS/MAPK pathway were identified via whole-exome sequencing (WES) in either of the two resistant cell lines after venetoclax selection. The addition of RMC-7977 re-sensitized both resistant cell lines to venetoclax (Figure 4C-D). Using dynamic iBH3 profiling, we determined that venR Molm-14 and Molm-14 *NRAS*^G12C^ cells have a lower apoptotic dependency on BCL2 (measured as cytochrome C release upon exposure to either BH3-mimetic inhibitors), but a higher dependency on MCL1 and, to a lesser extent, to BCLxL, compared to the parental cell lines (Figure 4E-F). Additionally, venR cells express higher levels of MCL1 protein (Figure 4G). These observations are consistent with previous *in vitro* findings in AML cell lines resistant to venetoclax.^30^ Treatment with RMC-7977 downregulated MCL1 in venR cells more profoundly than in the parental cell lines and induced a deeper suppression of ERK and AKT phosphorylation. We also observed an electrophoretic shift of BCLxL in venR cells treated with RMC-7977, suggestive of post-translational modifications, and a loss of full-length pro-apoptotic BID, likely due to cleavage (Figure 4G). Unexpectedly, in venR cells, RMC-7977 did not significantly increase the expression of pro-apoptotic proteins BIM and BMF, though it did so in the parental cells (Figure 4G). Upregulation of BIM and BMF has been previously observed in AML cell lines after treatment with MAPK pathway inhibitors.^62,63^ These findings suggest that RMC-7977 induces cell death via a distinct mechanism in cells with acquired resistance to venetoclax compared to parental cells, and that this mechanism is dependent on RAS signaling. Accordingly, we found that Molm-14 and Molm-14 *NRAS*^G12C^ venR cells are more sensitive to apoptosis triggered by RMC-7977 treatment, compared to parental cells (Figure 4H-I).

We and others have previously demonstrated that effective co-inhibition of the RAS/MAPK pathway and BCL2 has a strong synergistic activity in AML cell lines.^62–65^ Indeed, we found that RMC-7977 and venetoclax showed high *in vitro* synergy in cell viability assays (assessed by the Bliss independence model). This synergistic activity was most notable in cell lines with low baseline sensitivity to venetoclax monotherapy, driven by either RAS mutations or activating upstream RTK mutations (Figure 4J).

### RMC-7977 combinations are active in primary AML samples with RAS/MAPK-activating signaling mutations

To evaluate the activity of RMC-7977 and its combinations in primary AML samples, we performed colony-forming unit (CFU) assays using samples from three patients with AML harboring both *FLT3* and *NRAS* or *KRAS* mutations. Samples were treated with vehicle (DMSO), RMC-7977 (25nM), gilteritinib (50nM), venetoclax (500 nM) or combinations of RMC-7977 with gilteritinib or venetoclax. In contrast to gilteritinib and venetoclax single agents, RMC-7977 and RMC-7977 plus venetoclax significantly inhibited colony formation in all three samples, whereas RMC-7977 + gilteritinib did so in two out of the three samples (Figure 5A-C). We did not observe any reduction in number of colonies in two healthy donor samples (Figure 5D-E) with either RMC-7977 or its drug combinations, suggesting that these combinations may have low hematopoietic toxicity clinically.

**Figure 5.**
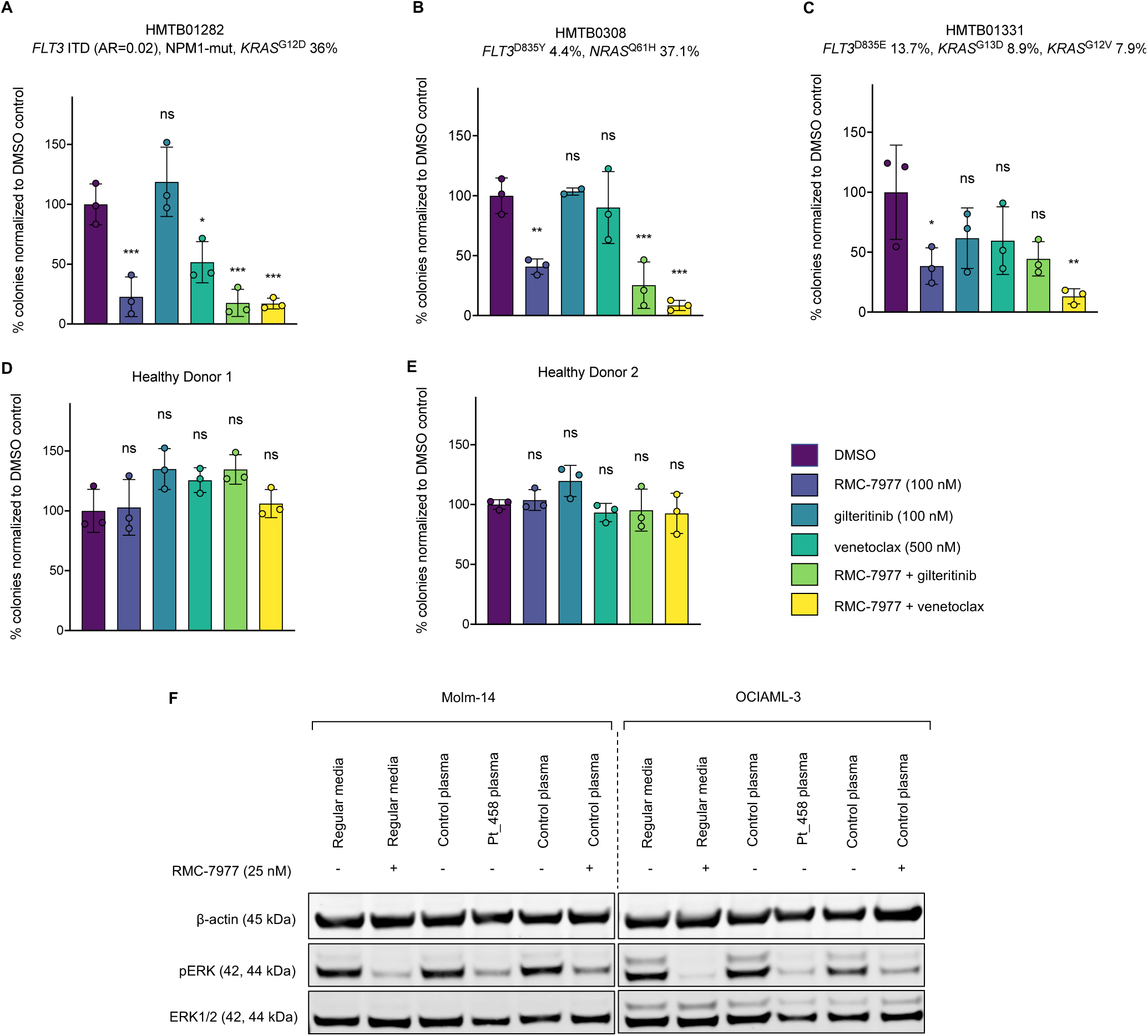
*Ex vivo* activity of RMC-7977 in primary AML samples. A-E: Differences in CFU-normalized counts in primary samples derived from individuals with AML with signaling mutations (A-C) or healthy donors (D-E). Data represent three replicates per sample; one-way ANOVA with Dunnet correction for multiple comparisons was used for statistical significance analysis (∗∗∗∗p ≤ 0.0001, ∗∗∗p ≤ 0.001, ∗∗p ≤ 0.01, ∗p ≤ 0.05, ns, non-significant). F: Western blot analysis of Molm-14 and OCIAML-3 cell lines exposed to either regular medium or plasma collected from a healthy donor in the presence or absence of RMC-7977 and cells exposed to plasma collected from a patient receiving daraxonrasib (RMC-6236, 200 mg daily).

Next, we sought to determine whether the plasma concentration of the clinical RAS(ON) inhibitor daraxonrasib (RMC-6236) reached in patients is sufficient to suppress RAS/MAPK signaling in AML cells as assessed by plasma inhibitory (PIA) assays. We incubated Molm-14 and OCIAML-3 cells for two hours in either plasma collected from a healthy donor or plasma collected at steady-state concentration from a patient receiving daraxonrasib (RMC-6236) 200 mg orally daily and pembrolizumab 200 mg intravenously every three weeks for NSCLC. The patient plasma inhibited ERK phosphorylation in both AML cell lines to an extent comparable to that seen with RMC-7977 at a concentration of 25 nM in regular culture media (Figure 5F). We also observed a substantial inhibition of phospho-ERK in cell lines incubated in control plasma treated with RMC-7977 at a concentration of 25 nM, suggesting that plasma protein binding could have minimal impact on target inhibition in leukemia cells. These *in vitro* data suggest potential clinical efficacy and therapeutic index, but will need clinical validation.

### RMC-7977 is active in therapeutic combinations in AML patient-derived xenografts

Given the promising *in vitro* activity of RMC-7977 in overcoming both FLT3i and BCL2i resistance, we next sought to assess the efficacy of RMC-7977 and its combinations *in vivo* using patient-derived xenograft (PDX) models. First, we performed a trial in a PDX model derived from a pediatric AML sample harboring *NRAS*^Q61L^ and a *KMT2A*::*PICALM* fusion (CPCT-0007, Figure 6A).^66,67^ Busulfan-conditioned NSG mice were injected, confirmed to have detectable engraftment by flow cytometry, and randomized into four treatment groups (n=6 mice/group), then treated orally with vehicle, venetoclax, RMC-7977, and the RMC-7977 plus venetoclax combination for 22 days. Throughout the trial, disease progression was monitored by the percentage of hCD45+ cells in peripheral blood via flow cytometry. At study termination or at humane endpoint, mice were sacrificed, and hCD45+ cells from cardiac blood, spleen, and bone marrow were measured via flow cytometry. We observed that while venetoclax had no activity as monotherapy in any tissue, RMC-7977 alone or in combination with venetoclax led to a complete suppression of hCD45+ cells in blood, spleen, and bone marrow (Figure 6B-D).

**Figure 6.**
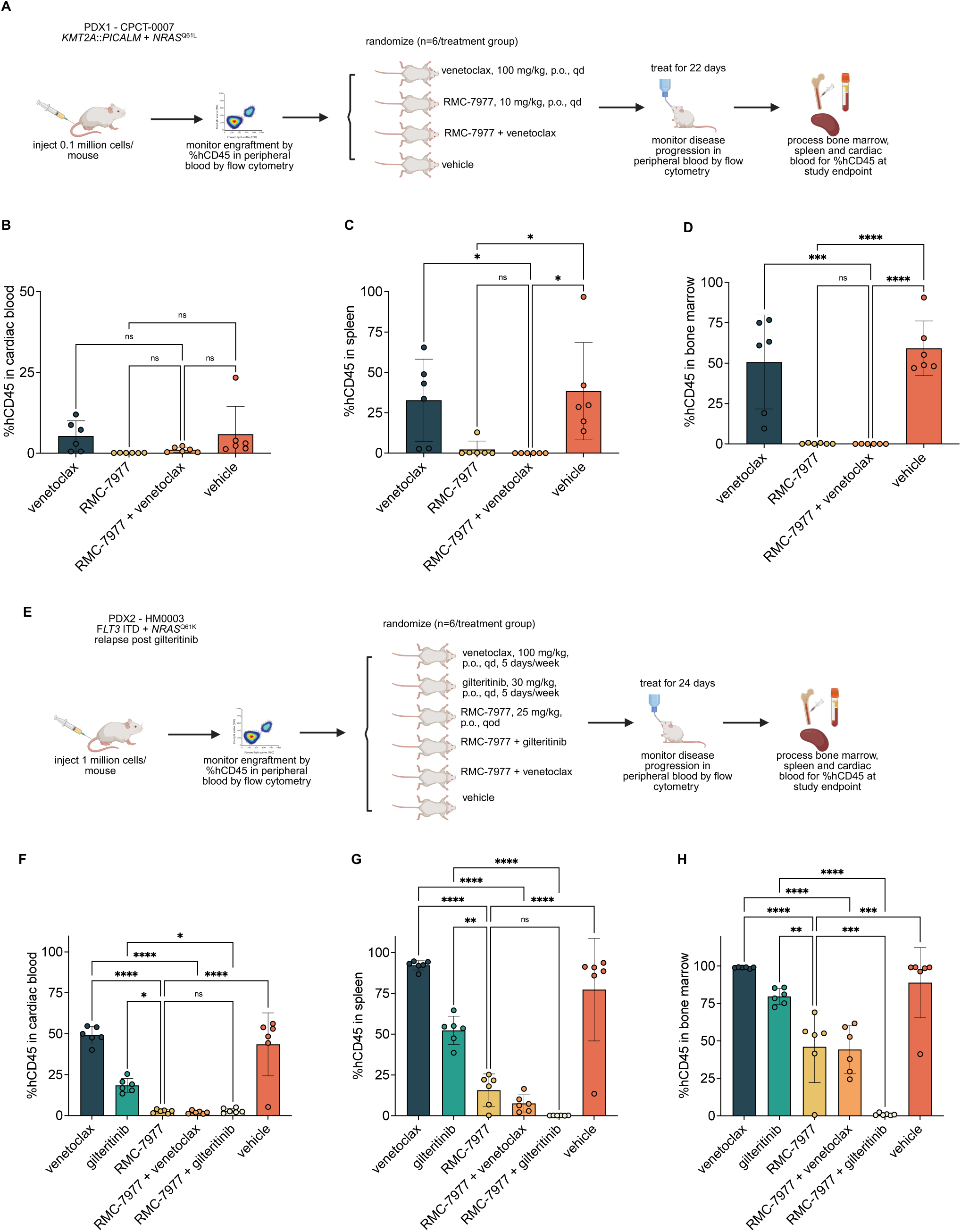
*In vivo* activity of RMC-7977 and therapeutic combinations. A and C: Schematic representations of PDX *in vivo* study designs. B-D and F-H: Quantification of hCD45+ cells in target organs for mice in each treatment group at study termination (n=6 mice/group for both trials). Data represent the mean percentage of hCD45+ cells ± SD; one-way ANOVA with Tukey correction for multiple comparisons was used for statistical significance analysis (∗∗∗∗p ≤ 0.0001, ∗∗∗p ≤ 0.001, ∗∗p ≤ 0.01, ∗p ≤ 0.05, ns, non-significant).

Next, to investigate the activity of RMC-7977 in a model of RAS-driven resistance to FLT3i, we established a PDX model (HM0003) from a patient with *FLT3*-ITD^+^ AML who relapsed with a secondary *NRAS*^Q61K^ mutation after long-term treatment with the FLT3i gilteritinib (Figure 6E). NSG mice were then injected, confirmed to have detectable engraftment by flow cytometry, and randomized into six treatment groups (n=6 mice/group) and treated for 24 days with: vehicle, gilteritinib, venetoclax, RMC-7977, and combinations of RMC-7977 with either gilteritinib or venetoclax. At study termination, mice that received RMC-7977 alone or in combination with either gilteritinib or venetoclax exhibited a significant reduction of leukemic burden across all tissues (Figure 6F-H). Strikingly, the combination of RMC-7977 and gilteritinib led to a profound antileukemic effect, even in the bone marrow (average hCD45+ of 0.5%), a tissue compartment where all other single agents and the combination of RMC-7977 and venetoclax failed to clear leukemic cells (Figure 6H). This observation is consistent with our *in vitro* findings that RAS(ON) inhibition re-sensitizes *FLT3* and RAS co-mutated AML to the antileukemic activity of FLT3i and highlights the translational potential of this therapeutic combination in relapsed *FLT3*^mut^ AML.

We did not observe any significant change in body weight between study groups in either of the trials. (Supplementary figures S5A-B). Mice receiving RMC-7977 either as monotherapy or in combinations had persistently lower percentages of circulating hCD45+ cells compared to those who were treated with vehicle, gilteritinib or venetoclax monotherapies in both trials (Supplementary figures S5B-C).

## Discussion

Mutations in *NRAS* or *KRAS* are frequent drivers of relapse and resistance to therapies targeting FLT3 and BCL2 in AML.^17–20,24,25^ Patients who relapse on targeted therapies lack effective treatments and have a dismal prognosis. Given the central role of RAS signaling in the biology of R/R AML, agents that effectively target RAS have the potential for significant clinical impact. Most direct RAS inhibitors currently in clinical development target specific *KRAS* mutations.^42,43,45,56^ In AML, however, therapeutic resistance often emerges with heterogenous, polyclonal RAS mutations (most frequently in *NRAS*), rather than with a single, recurrent hotspot mutation. Moreover, increased RAS activation associated with resistance can occur in RAS^WT^ clones.^17,21^ Therefore, we reason that broad-spectrum inhibition of both mutant and wild-type RAS isoforms is a suitable strategy to address therapy resistance in AML. In this study, we have shown that the RAS(ON) multi-selective inhibitor RMC-7977 exhibits potent anti-leukemic activity in preclinical models of FLT3i-resistant and BCL2i-resistant AML driven by activated RAS signaling.

Previous attempts to therapeutically target the MAPK pathway in AML have focused on downstream pathway effector inhibition, generally without success. Early-phase trials of MEKi showed only modest response rates as monotherapy,^35–38^ while combination regimens with MEKi and venetoclax, HMA plus venetoclax, or PI3K inhibitors did not show significant clinical benefit and suffered from excess toxicity.^68–70^ This lack of benefit may be due to the fact that MEKi are cytostatic rather than pro-apoptotic, or due to their vulnerability to feedback signaling reactivation.^71,72^ Pan-RAF inhibitors may overcome these limitations to some extent, but they are not validated clinically in AML.^72–75^ We have shown that RMC-7977 can both arrest cell proliferation and reliably induce apoptosis in AML cells with activated RAS signaling *in vitro*, suggesting that it may have improved clinical anti-leukemic properties compared to downstream MAPK inhibitors. This effect may be in part attributable to disruption of RAS-PI3K p110α interaction, as expression of constitutively active PI3K mutants (*PIK3CA*^H1047R^ and *PIK3CA*^E542K^) attenuated the apoptotic response to RMC-7977. In *FLT3*-mut AML, signaling through both MAPK and PI3K pathways is simultaneously activated through upstream RTK activity, but PI3K signaling alone is not sufficient to cause resistance to FLT3i.^80^ However, we found that in combination with gilteritinib, RMC-7977 leads to a more profound suppression of PI3K signaling compared to gilteritinib alone in *FLT3* and *NRAS* co-mutated cells with higher PI3K output attributable to hyperactive RAS signaling.

Resistance to FLT3 inhibition is an important clinical problem in AML and can be caused by selection for off-target mutations in downstream signaling effectors, most commonly RAS, or on-target mutations in *FLT3* itself.^16^ Our data suggest that RMC-7977 may be helpful in overcoming both types of resistance. We modeled the clonal competition and selection for RAS mutations that often leads to clinical gilteritinib resistance using *in vitro* co-culture experiments of cell lines harboring *FLT3*-ITD or *FLT3*-ITD plus *NRAS*^Q61K^, which showed that the addition of RMC-7977 to gilteritinib abolishes the fitness advantage of RAS-mutant cells. We also observed profound anti-leukemic activity in a PDX model derived from a patient who relapsed on gilteritinib with an acquired *NRAS*^Q61K^ mutation. The combination of FLT3 and RAS inhibition achieved near-complete eradication of leukemic blasts in peripheral blood, spleen, and bone marrow of mice, in contrast to gilteritinib monotherapy. These findings suggest that RMC-7977 may have a role in preventing or reversing RAS-mediated resistance to gilteritinib. We also showed that RMC-7977 may overcome resistance to type II FLT3 inhibitors caused by on-target mutations rather than downstream signaling mutations.^19,58^ RMC-7977 retained activity in Molm-14 cells harboring both *FLT3*-ITD and TKD mutations, including the non-canonical “gatekeeper” *FLT3*^F691L^ mutation, associated with cross-resistance to gilteritinib.^77^ These data highlight the potential of direct RAS inhibition to broadly address FLT3 inhibitor resistance.

Hyperactivation of the RAS/MAPK pathway similarly mediates resistance to the BCL2 inhibitor venetoclax. RAS mutations have been identified at relapse on venetoclax in combinations with either HMA or gilteritinib.^21,24^ Moreover, *in vitro* studies suggest that RAS^mut^ LSCs preferentially give rise to AML with monocytic phenotype.^33^ Monocytic differentiation has been associated in AML with lower sensitivity to venetoclax and a higher anti-apoptotic dependence on MCL1, along with higher expression of *BCL2A1*.^21–23,78^ Using data from the BEAT AML 2.0 trial, we confirmed that a RAS transcriptional signature directly correlates with venetoclax resistance, monocytic differentiation, and elevated *BCL2A1* expression and inversely correlates with the *BCL2*/*MCL1* expression ratio. Intriguingly, we found that these associations persist even in samples without any detectable RAS mutations, suggesting that activated RAS signaling and its transcriptional consequences can drive a venetoclax-resistant monocytic phenotype even in the absence of RAS mutation *per se*. This finding is supported by results from longitudinal single-cell multiomic sequencing studies of patients treated with gilteritinib and venetoclax.^21^ Accordingly, we hypothesized that broad inhibition of RAS(ON), both wild-type and mutant, can overcome this mechanism of resistance. We generated venetoclax-resistant cell lines in both RAS wild-type and RAS mutant genetic backgrounds and demonstrated that RMC-7977 re-sensitized them equally to venetoclax. These venetoclax-resistant cell lines have a higher expression and dependency on MCL1, in agreement with previous reports of similar biochemical changes in venetoclax-resistant AML cell lines.^30^ Consistently, we noticed that RMC-7977 induced a deeper apoptotic response and MCL1 downregulation in venetoclax-resistant than in venetoclax-sensitive cells, indicating a heightened addiction to the RAS/MAPK-MCL1 axis independent of RAS genotype, a vulnerability exploitable by RAS(ON) inhibition. These results underscore that RAS(ON) inhibition is a promising strategy to address venetoclax resistance in AML, an area of great unmet clinical need. Our findings further suggest that monocytic differentiation, a troublesome disease subtype often associated with venetoclax resistance in AML, may serve as a biomarker of RAS signaling activation and may be uniquely vulnerable to RAS(ON) inhibition. Further studies should explore the clinical potential of targeting RAS in combination with venetoclax in monocytic AML with and without RAS mutations.

Concomitant co-inhibition of the RAS/MAPK pathway and BCL2 has shown preclinical synergy in AML.^62,64,65^ Similarly, RMC 7977 demonstrated robust synergy with venetoclax in multiple cell lines harboring activating MAPK mutations. Additionally, in a venetoclax-resistant PDX model with a *KMT2A*::*PICALM* fusion and *NRAS*^Q61L^, RMC-7977 showed significant anti-leukemic activity, both as monotherapy and in combination with venetoclax. In this model, we were unable to discern any significant difference in response between RMC-7977 monotherapy and the combination therapy, likely due to the high sensitivity of this PDX to RAS(ON) inhibition alone. These data highlight the therapeutic potential of RAS(ON) inhibition to target, alone or in combination with venetoclax, RAS-mutant leukemia.

A potential concern for combining broad spectrum RAS inhibition with other targeted agents in AML is tolerability. By virtue of their selectivity for active, GTP-bound RAS conformation, RAS(ON) inhibitors preferentially target cells with a higher addiction to MAPK signaling.^46,52^ Daraxonrasib has shown a manageable safety profile in solid malignancies trials, with overall mild hematologic toxicitiy.^48,49^ In our PDX trials, RMC-7977 (alone or in combination with gilteritinib or venetoclax) did not induce significant weight loss or overt toxicity, nor did it impair CFU formation from healthy donor bone marrow. These studies suggest a therapeutic window for RAS(ON) inhibition in AML, but safety and optimal dosing in AML will need to be determined clinically.

As with other targeted therapies, resistance to RAS(ON) inhibition is anticipated. Our finding that activating downstream mutations, either in PI3K/AKT or MEK, can confer resistance to apoptosis induced by RAS(ON) inhibition, suggests these as potential mechanisms of resistance, though such mutations are relatively uncommon in AML. Preclinical data implicate copy number gains of *MYC* or compensatory activation of WNT, NF-κB or JAK/STAT pathways as potential adaptive responses and escape mechanisms.^52,56^ Of note, we observed increased JAK/STAT pathway activity scores in AML cell lines treated with RMC-7977 via RNA-seq and PROGENy analysis. Future studies should comprehensively characterize escape mechanisms and inform on rational therapeutic strategies to mitigate resistance to RAS(ON) inhibitors in AML.

In conclusion, our study provides strong preclinical evidence that broad-spectrum, multi-selective RAS(ON) inhibition is a feasible therapeutic strategy to prevent and overcome RAS/MAPK-mediated resistance to FLT3 and BCL2 inhibitors in AML. Given the critical role of resistance in limiting the long-term efficacy of currently approved targeted therapies, clinical trials evaluating daraxonrasib in combination with either FLT3 or BCL2 inhibitors are urgently warranted to improve outcomes of patients with R/R AML.

## Methods

### Cell lines

Molm-14, MV4-11, OCIAML-3, HL-60, and NOMO-1 were a gift from Dr. Scott Kogan (University of California, San Francisco, San Francisco, CA). Kasumi-1 and SKNO-1 cell lines were purchased from ATCC. Molm-14 *NRAS*-mutant (*NRAS*^G12C^ or *NRAS*^Q61K^) and *FLT3* TKD–mutant cells were generated by culturing Molm-14 parental cells *in vitro* in media containing escalating doses of quizartinib (0.5–20 nmol/L).^79^ Quizartinib-resistant cells were subcloned and genotyped by Sanger sequencing. Venetoclax-resistant (venR) cells were generated by culturing Molm-14 or Molm-14 *NRAS*^G12C^ cells in escalating doses of venetoclax (155-6000 nmol/L). To generate Molm-14 cell lines with doxycycline-inducible overexpression of *PIK3CA*, *AKT1*^E17K^, MEK^DD^ or *NRAS* mutations, the respective mutant variants were cloned into a Gateway tetracycline-inducible destination lentiviral vector, pCW57.1 (Addgene cat #41393). Cloning was performed by Twist Biosciences. Lenti-X 293T cells (Takara Bio) were infected using Lipofectamine 3000 (Invitrogen) and cultured for 48 hours in DMEM with 10% FBS. The supernatant was harvested, concentrated using Lenti-X concentrator (Takara Bio), then used to infect cell lines. Forty-eight hours following lentiviral infection, cells were selected with puromycin. Molm-14 cells expressing mCherry and Molm-14 *NRAS*^Q61K^ cells expressing Zs.Green fluorescent protein were a gift from Neil Shah (University of California, San Francisco, San Francisco, CA). Cell lines were cultured in RPMI-1640 (Gibco) with 10% fetal bovine serum (FBS) and 1% penicillin/streptomycin/L-glutamine (Gibco), with the exception of Kasumi-1 and HL-60 cells, cultured with 20% FBS. Culture media for SKNO-1 cell line was supplemented with human GM-CSF (10 ng/mL, PeproTech). All cells tested negative for Mycoplasma by the MycoAlert PLUS Mycoplasma Detection Kit (Lonza). Experiments were performed within three months of cell line thawing. Authentication of all cell lines was performed at the University of California, Berkeley, DNA Sequencing Facility, using short tandem repeat (STR) DNA profiling.

### Patient samples

Collection, storage, and experimental studies of de-identified primary AML samples, plasma or hematopoietic stem cells from healthy donors were conducted after approval of institutional review board via tissue banking protocols from the University of California San Francisco (Hematologic Malignancies Tissue Bank and Molecular Profiling in Thoracic Malignancies). Informed, written consent according to the Declaration of Helsinki was obtained from patients prior to tissue collection.

### Compounds

RMC-7977 was provided by Revolution Medicines. Gilteritinib (S7754), quizartinib (S1526), naporafenib (S8745), trametinib (S2673), ulixertinib (S7854), venetoclax (S8048), WEHI-539 (S7100) and were purchased from Selleckchem. S63845 (HY-100741) was purchased from MedChem Express. BIM peptide was purchased from New England Peptide.

### Cell viability assays

Cells were seeded at a concentration of 0.2e^6^/ml in three technical replicates in 96-well plates and exposed to increasing concentrations of the indicated inhibitors for 48 hours. For doxycycline-inducible cell line experiments, cells were stimulated for 24 hours with doxycycline 1 μg/mL and maintained in the same concentration of doxycycline for the duration of experiments. Cell viability was assessed using CellTiter-Glo Luminescent Cell Viability Assay (Promega), and luminescence was measured on a Molecular Devices iD3 multimode plate reader and normalized to untreated controls.

### Apoptosis assays

4 x10^4^ cells were seeded in three technical replicates in 96-well clear plates and exposed to increasing concentrations of indicated drug for 24 hours. CellEvent Caspase-3/7 Green Flow Cytometry Kit (Thermo Fisher) was used to measure fraction of apoptotic cells by flow cytometry (Agilent NovoCyte 3005 with NovoSampler Pro plate adapter or BD LSR Fortessa with HTS sampler). Percentage of caspase-3/7 negative cells in each treatment condition was normalized to vehicle-treated control populations (NovoExpress, Agilent or FlowJo software v9, BD Biosciences).

### Western Blot

Cells were plated in appropriate media and treated with the indicated concentrations of inhibitors. After 2 hours or 24 hours incubation time, cells were washed in PBS and lysed in buffer (50 mmol/L HEPES, pH 7.4, 10% glycerol, 150 mmol/L NaCl, 1% Triton X-100, 1 mmol/L EDTA, 1 mmol/L EGTA, and 1.5 mmol/L MgCl_2_) supplemented with protease and phosphatase inhibitors (EMD Millipore). The lysates were clarified by centrifugation, quantitated by BCA assay (Thermo Scientific), and normalized. 20 mg of protein were loaded on 10% Bis-Tris gels and then transferred to nitrocellulose membranes. Immunoblotting was performed using the following antibodies from Cell Signaling Technology: anti-β-Actin (8H10D10, #3700), anti-Erk1/2 (3A7, #9107), anti-phoshpo-Erk1/2 (Thr202/Tyr204, #9101), anti-cRAF (#9422), anti-phospho-cRAF (Ser338, 56A6, #9427), anti-MEK (61B12, #2352), anti-phosphoMEK1/2 (Ser217/221, #9121), anti-RSK1/RSK2/RSK3 (32D7, #9355), anti-phospho-p90RSK (Thr359/Ser363, #9344), anti-Akt (#9272), anti-phospho Akt (Thr308, #9275), anti-S6 ribosomal protein (54D2, #2317), anti-phospho-S6 ribosomal protein (Ser235/236, #2211), anti-Stat5 (D2O6Y, #94205), anti-phospho-Stat5 (Tyr 694, #9351), anti-Bcl2 (124, #15071), anti-Bcl-xL (#2762), anti-Mcl-1 (#4572), anti-Bim (C34C5, #2933), anti-BMF (E5U2J, #50542), anti-BID (3C5, #8762), anti-BAD (#9292). Nitrocellulose membranes were subsequently incubated in a solution of secondary antibodies (IRDye 800CW Goat anti-Rabbit, #935-32211, IRDye 680RD Goat anti-Mouse, #935-68070, LI-COR Biosciences) and scanned on an Odyssey CLx infrared imaging system (LI-COR Biosciences).

### Bulk RNA sequencing

Cells were treated for either 2 or 24 hours, with three to four biological replicates per condition. Messenger RNA (mRNA) was purified from total RNA using poly-T oligo-attached magnetic beads. All RNA samples exhibited RIN scores ≥8.5. Following fragmentation, first-strand cDNA was synthesized using random hexamer primers, followed by second-strand synthesis. Library preparation included end repair, A-tailing, adapter ligation, size selection, amplification, and purification. Libraries were quantified using Qubit and real-time PCR, and fragment size distribution was assessed using a Bioanalyzer. Quantified libraries were pooled and sequenced on an Illumina platform using a 2×150 bp paired-end configuration. Raw FASTQ files were processed through a Snakemake pipeline (v9.3.1), which incorporated quality control assessments using MultiQC (v1.28). Adapter sequences and low-quality bases were removed using Trimmomatic (v0.36). Cleaned reads were aligned to the human reference genome (hg38) using STAR (v2.7.8a) and quantified with RSEM (v1.3.3). Normalization of aligned read counts was performed using the trimmed mean of M-values (TMM) method implemented in the EdgeR package (v3.24.3), followed by log2 transformation of the counts. The mean-variance relationship was modeled using the VOOM function from the limma package (v3.38.3) to assign precision weights and perform differential expression analysis. Genes were annotated using the packages STRING (v.12.0.0) and panther (v. 1.0.12) databases on R. Gene set enrichment analysis (GSEA v4.3.3) and clusterProfiler (v4.16) were subsequently employed to identify significantly enriched functional gene sets. The decoupleR package^80^ was used to perform the PROGENy analysis to infer signaling activity of pathways from gene expression.^81^

### Clonal outgrowth assay

Fluorescently-tagged Molm-14 (mCherry) cells and Molm-14 *NRAS*^Q61K^ (Zs.Green) cells were mixed in a ratio 1:1 in three replicates per condition, then incubated for 96 hours in the presence of DMSO, RMC-7977, gilteritinib or RMC-7977 + gilteritinib. After 96 hours, cell mixtures were stained with a live/dead stain, Sytox AADvanced (Thermo Fisher, MA, USA), then counted on a flow cytometer (BD LSR Fortessa with HTS sampler). Data were analyzed using the FlowJo software v9 (BD Biosciences).

### BEAT AML data mining

RNA sequencing (normalized expression), whole exome sequencing/targeted sequencing mutation data and *ex vivo* drug sensitivity from the BEAT AML2.0 trial were downloaded from https://biodev.github.io/BeatAML2/. The single-sample GSEA^82^ (ssGSEA) module of GenePattern^83^ was performed to determine individual sample enrichment scores.

### Dynamic BH3 profiling

The BH3 profiling of AML cell lines was carried out as previously described.^84^ In brief, cells were incubated in 96-well plates in a solution of MEB buffer containing 0.002% digitonin and either Bim peptide or BCL, BCLxL or MCL1 inhibitors, fixed and stained with an anti-cytochrome C (clone 6H2.B4, # 612310, BioLegend). Flow cytometry analysis was performed on an Agilent NovoCyte 3005 instrument. Cytochrome C release was measured by measuring the cytochrome C median fluorescent intensity (MFI), normalized to vehicle-treated control populations (NovoExpress, Agilent).

### Patient-derived xenograft models

All mouse experiments were conducted in accordance with an approved protocol by the UCSF Institutional Animal Care and Usage Committee. Patient derived xenograft models of human AML were generated in NOD-scid IL2Rγ^null^ mice (NSG, Jackson Laboratory) and serially passaged. Mice were then injected with 0.1 to 1 million PDX cells. Mice were monitored for weight change weekly during the entirety of the trial. To confirm engraftment, mice were bled by submandibular puncture weekly and analyzed by flow cytometry for human (hCD45) and mouse CD45 (mCD45). Mice were randomly assigned to a treatment arm once they reached greater than or equal to (0.5% hCD45+ cells / total events) by flow cytometry. For the PDX1 trial (CPCT-0007), the treatment arms were defined as following: vehicle (oral gavage, daily), RMC-7977 (10 mg/kg, oral gavage, daily), venetoclax (100 mg/kg, oral gavage, daily), RMC-7977 (10 mg/kg, oral gavage, daily) plus venetoclax (100 mg/kg, oral gavage, daily). For the PDX2 trial (HM-0003), the treatment arms were defined as following: vehicle (oral gavage, daily, five days per week), RMC-7977 (25 mg/kg, oral gavage, every other day), gilteritinib (30 mg/kg, daily, five days per week), venetoclax (100 mg/kg, oral gavage, five days per week), RMC-7977 (25 mg/kg, oral gavage, every other day) plus gilteritinib (30 mg/kg, daily, five days per week) and RMC-7977 (25 mg/kg, oral gavage, every other day) plus venetoclax (100 mg/kg, oral gavage, five days per week). Mice were treated for 22 and 24 days, respectively, during which they were monitored daily for symptomatic disease progression and every other week for disease burden by flow cytometry. Mice were euthanized after 28 days or when they reached humane endpoint, whichever was sooner. Mice were processed upon endpoint, and cardiac blood, bone marrow, and spleen were analyzed by flow cytometry for hCD45+/mCD45+.

### Colony-forming unit assays

Primary samples from leukapheresis products harvested from either healthy individuals or patients with AML were thawed in DMEM media (Gibco) supplemented with 20% FBS, 2 mM EDTA and 500 μg DNase I. 2 x 10^5^ cells per treatment condition, in three replicates, were incubated with DMSO, RMC-7977 (100 nM), gilteritinib (100 nM), venetoclax (500 nM) or the combination of RMC-7977 with either inhibitor in methylcellulose media enriched with human cytokines (recombinant human SCF, GM-CSF, G-CSF, IL-3, IL-6, Epo, #HSC005, R&D Systems). The methylcellulose mixture was plated in three replicates to 35 mm culture dishes using regular syringes with 16-gauge non-stick needles and incubated at 37°C, 5% CO2 for 14-16 days. Colony-forming units were counted using an Olympus CKX53 light microscope with a 4X magnification objective, and the counts were normalized to untreated controls.

### Plasma inhibitory assays

Molm-14 and OCIAML-3 cell lines were incubated for 24 hours in either plasma collected from a healthy human donor or plasma collected at steady state concentration from one patient receiving daraxonrasib (200 mg orally, daily) and pembrolizumab (200 mg intravenously, every three weeks) within the NCT06162221 (Study of RAS(ON) Inhibitor Combinations in Patients with Advanced RAS-mutated NSCLC) clinical trial. For control conditions, same cells were incubated in either RPMI 10%, RPMI 10% + RMC-7977 (25 nM) or healthy plasma + RMC-7977 (25 nM). After inhibitor incubation, cells were lysed and a western blot (See Western Blot methods above) was performed to assess ERK, AKT and ribosomal S6 phosphorylation in each condition. β-actin was used as a loading control.

### Statistical analysis

Details on statistical analysis of experiments can be found in the figure legends. Unpaired t tests or unpaired Mann-Whitney tests were used to compare two independent groups. One-way or two-way analysis of variance (ANOVA) tests were used to compare more than two groups, and correction for multiple comparisons was applied when needed. Statistical significance is denoted as follows: ∗, p ≤ 0.05; ∗∗, p ≤ 0.01; ∗∗∗, p ≤ 0.001; ∗∗∗∗, p ≤ 0.0001, unless stated otherwise. Data are reported as mean ± SD, unless otherwise specified. For cell viability readouts, dose-response curves and absolute IC_50_ values were calculated using a four-parameter nonlinear regression model. Drug synergy assessment was performed using the Bliss method within SynergyFinder v3.0 (https://synergyfinder.fimm.fi/), with LL4 curve fitting and cell viability readout data. GraphPad Prism software (v10) was used for statistical analysis and data plotting. BioRender.com was used to draw graphic illustrations.

## Data availability

RNA-sequencing data reported in this study have been deposited at the National Center for Biotechnology Information Gene Expression Omnibus repository and the accession number was pending at the time of this submission.

## Acknowledgements

The authors thank Revolution Medicines for providing RMC-7977, the Smith lab members, and Bianca Lee for scientific input and technical support throughout the project, Rashi Ragulan and Ximo Pechuan-Jorge for assistance with bioinformatics analysis, and John Byrd for constructive scientific discussions. This work was supported in part by Revolution Medicines and R01CA277031 (to C.C.S). C.A.C.P. was funded by a Young Investigator Award from the Alex’s Lemonade Stand Foundation. C.C.S. is a Leukemia & Lymphoma Society Scholar in Clinical Research and a Damon Runyon-Richard Lumsden Foundation Clinical Investigator supported (in part) by the Damon Runyon Cancer Research Foundation (CI-99-18).

## Authorship Contributions

Conceptualization, B.P. and C.C.S.; methodology, B.P., M.F.J., C.A.C.P., E.S. and C.C.S.; software, B.P. and N.M.; validation, B.P., M.F.J., M.P., E.T., A.K. and I.L.; formal analysis, B.P., M.F.J., E.T., A.K., I.L., N.M. and S.A.; investigation, B.P., M.F.J., M.P., E.T., A.K., I.L., S.A., C.M. and J.R.; resources, Y.P., M.L.C., A.C.L., E.S., and C.C.S.; data curation, B.P., M.F.J., E.T., A.K., I.L., C.A.C.P, N.M. and C.C.S.; writing – original draft, B.P.; writing – review & editing, B.P., M.F.J, C.A.C.P., Y.P., A.C.L., E.S. and C.C.S.; visualization, B.P., M.P., E.T., A.K. and N.M; supervision, E.S. and C.C.S.; project administration, B.P. and C.C.S.; funding acquisition, C.C.S.

## Statement of Prior Presentation

Presented partly in abstract form at the 65^th^ and 66^th^ annual meetings of the American Society of Hematology, San Diego, CA, 10 December 2023 and 8 December 2024.

