## Supplementary data for "Multi-selective RAS(ON) Inhibition Targets Oncogenic RAS Mutations and Overcomes RAS/MAPK-Mediated Resistance to FLT3 and BCL2 Inhibitors in Acute Myeloid Leukemia"

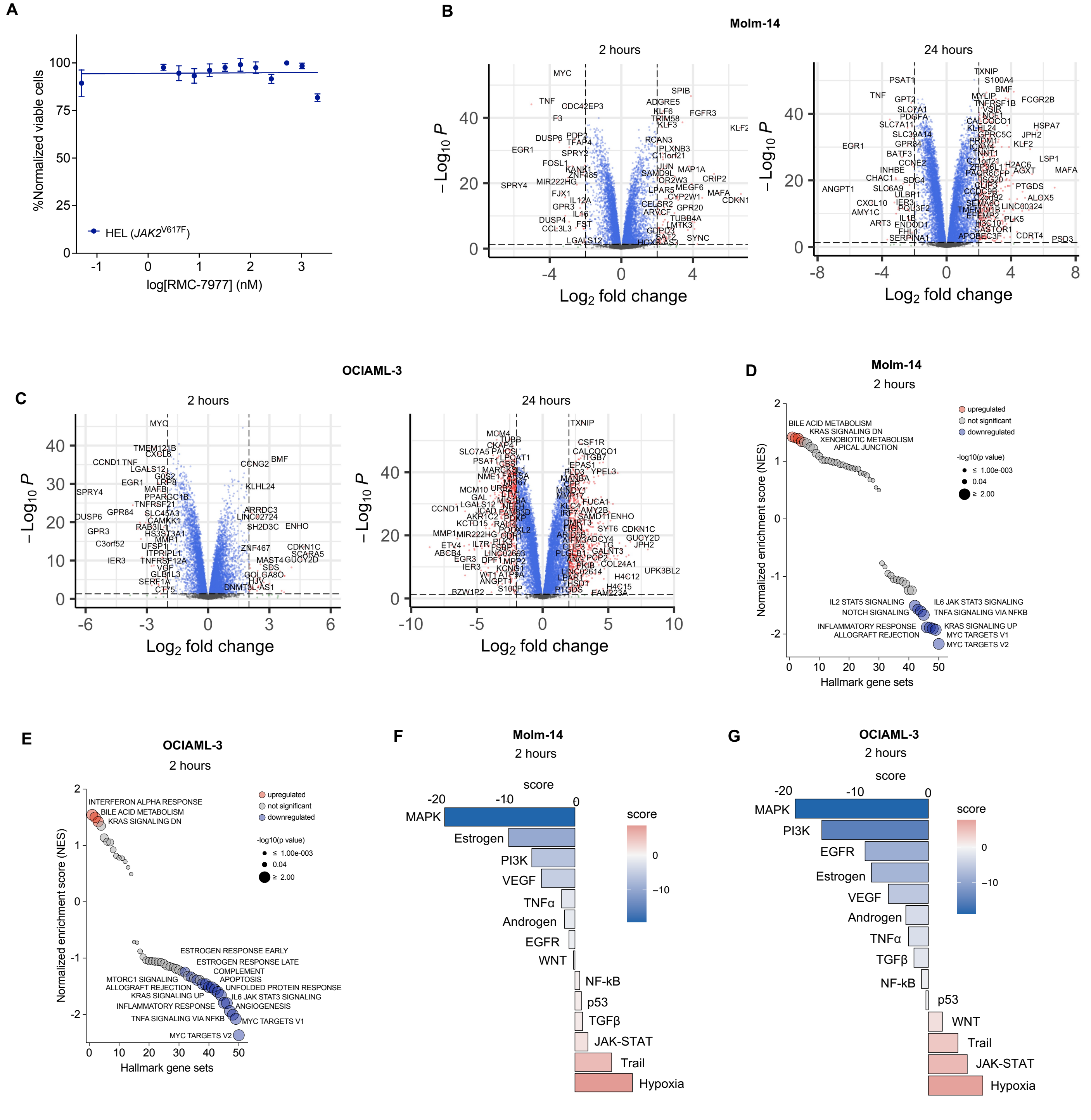

A
%Normalized viable cells
HEL (JAK2V617F)
log[RMC-7977] (nM)
B
Molm-14
2 hours
24 hours
−Log10 P
Log2 fold change
C
OCIAML-3
2 hours
24 hours
−Log10 P
Log2 fold change
D
Molm-14
2 hours
upregulated
not significant
downregulated
-log10(p value)
≤ 1.00e-003
0.04
≥ 2.00
BILE ACID METABOLISM
KRAS SIGNALING DN
XENOBIOTIC METABOLISM
APICAL JUNCTION
IL2 STAT5 SIGNALING
NOTCH SIGNALING
IL6 JAK STAT3 SIGNALING
TNFA SIGNALING VIA NFKB
INFLAMMATORY RESPONSE
ALLOGRAFT REJECTION
KRAS SIGNALING UP
MYC TARGETS V1
MYC TARGETS V2
Normalized enrichment score (NES)
Hallmark gene sets
E
OCIAML-3
2 hours
upregulated
not significant
downregulated
-log10(p value)
≤ 1.00e-003
0.04
≥ 2.00
INTERFERON ALPHA RESPONSE
BILE ACID METABOLISM
KRAS SIGNALING DN
ESTROGEN RESPONSE EARLY
ESTROGEN RESPONSE LATE
COMPLEMENT
MTORC1 SIGNALING
APOPTOSIS
ALLOGRAFT REJECTION
UNFOLDED PROTEIN RESPONSE
KRAS SIGNALING UP
IL6 JAK STAT3 SIGNALING
INFLAMMATORY RESPONSE
ANGIOGENESIS
TNFA SIGNALING VIA NFKB
MYC TARGETS V1
MYC TARGETS V2
Normalized enrichment score (NES)
Hallmark gene sets
F
Molm-14
2 hours
score
MAPK
Estrogen
PI3K
VEGF
TNFα
Androgen
EGFR
WNT
NF-kB
p53
TGFβ
JAK-STAT
Trail
Hypoxia
G
OCIAML-3
2 hours
score
MAPK
PI3K
EGFR
Estrogen
VEGF
Androgen
TNFα
TGFβ
NF-kB
p53
WNT
Trail
JAK-STAT
Hypoxia

**Supplementary Figure S1 – Activity of RAS(ON) multi-selective inhibitor RMC-7977 in AML cell lines.** A: Concentration-response curves representing relative proliferation of HEL cell line driven by *JAK2*$^{V617F}$ mutation after 48 h of exposure to serial doses of RMC-7977. Data represent the mean ± SD of three replicates. B-C: Volcano plots of differentially expressed genes in Molm-14 (B) and OCIAML-3 (C) cells after treatment with RMC-7977 for 2 hours and 24 hours respectively. Highlighted genes with a |log2FC|≥2 with a significance threshold of p value < 0.05. D-E: Bubble plots showing enrichment of GSEA Hallmark gene sets expression in Molm-14 and OCIAML-3 cell lines after exposure for 2 hours to RMC-7977 (25 nM). The size of the bubbles represents statistical significance, and the colors indicate upregulated (red) or downregulated (blue) expression. F-G: PROGENy pathway activity scores in Molm-14 and OCIAML-3 cells after exposure for for 2 hours to RMC-7977 (25 nM).

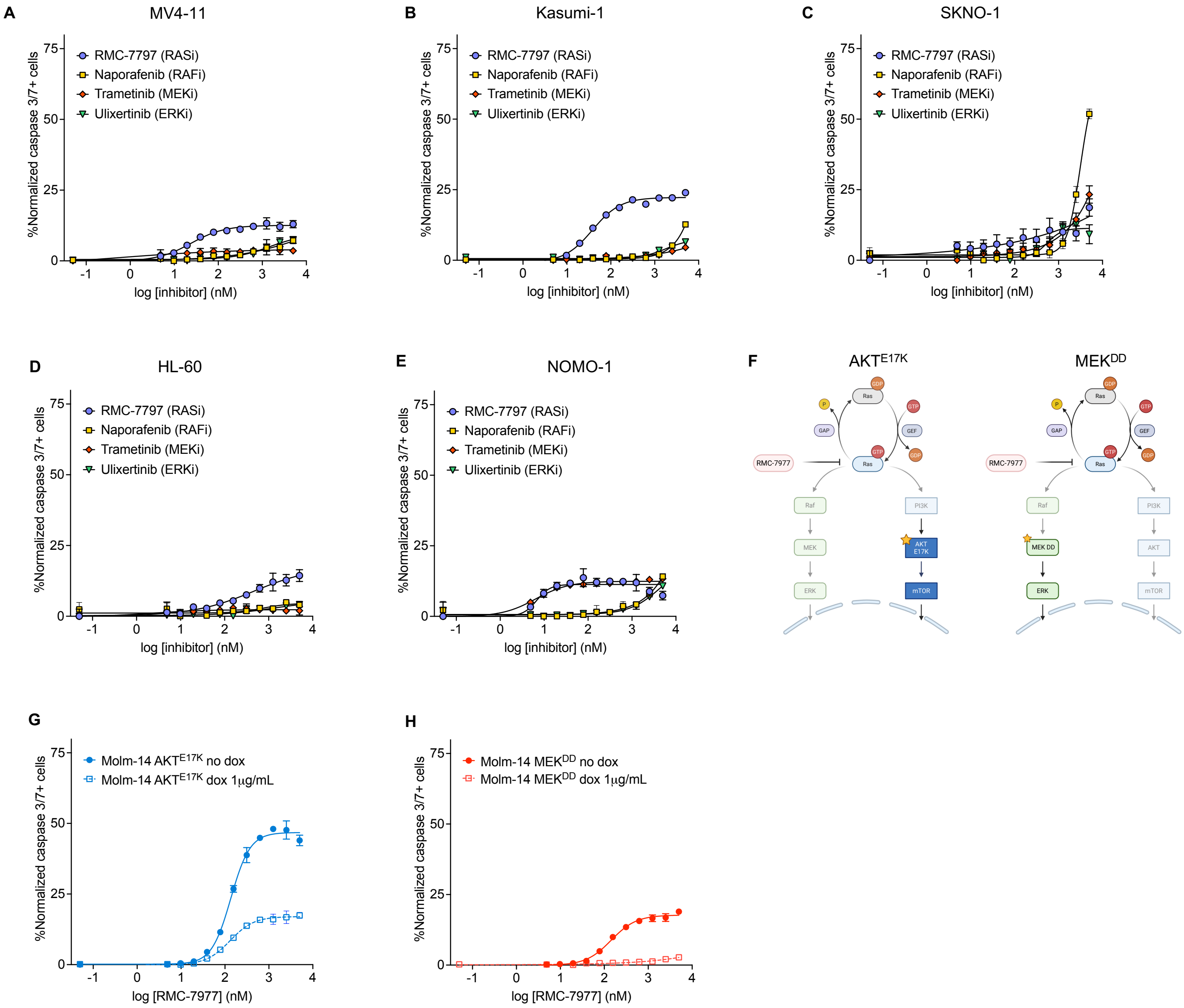
A
MV4-11
B
Kasumi-1
C
SKNO-1
D
HL-60
E
NOMO-1
%Normalized caspase 3/7+ cells
RMC-7797 (RASi)
Naporafenib (RAFi)
Trametinib (MEKi)
Ulixertinib (ERKi)
log [inhibitor] (nM)
F
AKT^E17K
MEK^DD
Ras
GDP
GTP
P
GAP
GEF
RMC-7977
Raf
MEK
ERK
PI3K
AKT E17K
mTOR
MEK DD
AKT
G
H
Molm-14 AKT^E17K no dox
Molm-14 AKT^E17K dox 1μg/mL
Molm-14 MEK^DD no dox
Molm-14 MEK^DD dox 1μg/mL
log [RMC-7977] (nM)

**Supplementary Figure S2 – RAS(ON) inhibition has a higher pro-apoptotic activity than downstream MAPK inhibition in AML cell lines.** A-E: Concentration-response curves showing activated caspase 3/7 expression in AML cell lines after 24 h of exposure to increasing doses of RMC-7977 (RASi), naporafenib (RAFi), trametinib (MEKi) and ulixertinib (ERKi), normalized to untreated controls. Data represent the means ± SD of three replicates. F: Schematic representation of constitutive active $AKT^{E17K}$ and $MEK^{DD}$ mutations. G-H: Comparative activated caspase 3/7 expression in Molm-14 cells transfected with doxycycline-inducible constructs overexpressing $AKT^{E17K}$ (F) or $MEK^{DD}$ after 24 hours of exposure to RMC-7977. Expression of AKT or MEK variants was induced with 1µg/mL of doxycycline 24 hours prior to inhibitor treatment. Data represent the means ± SD of three replicates.

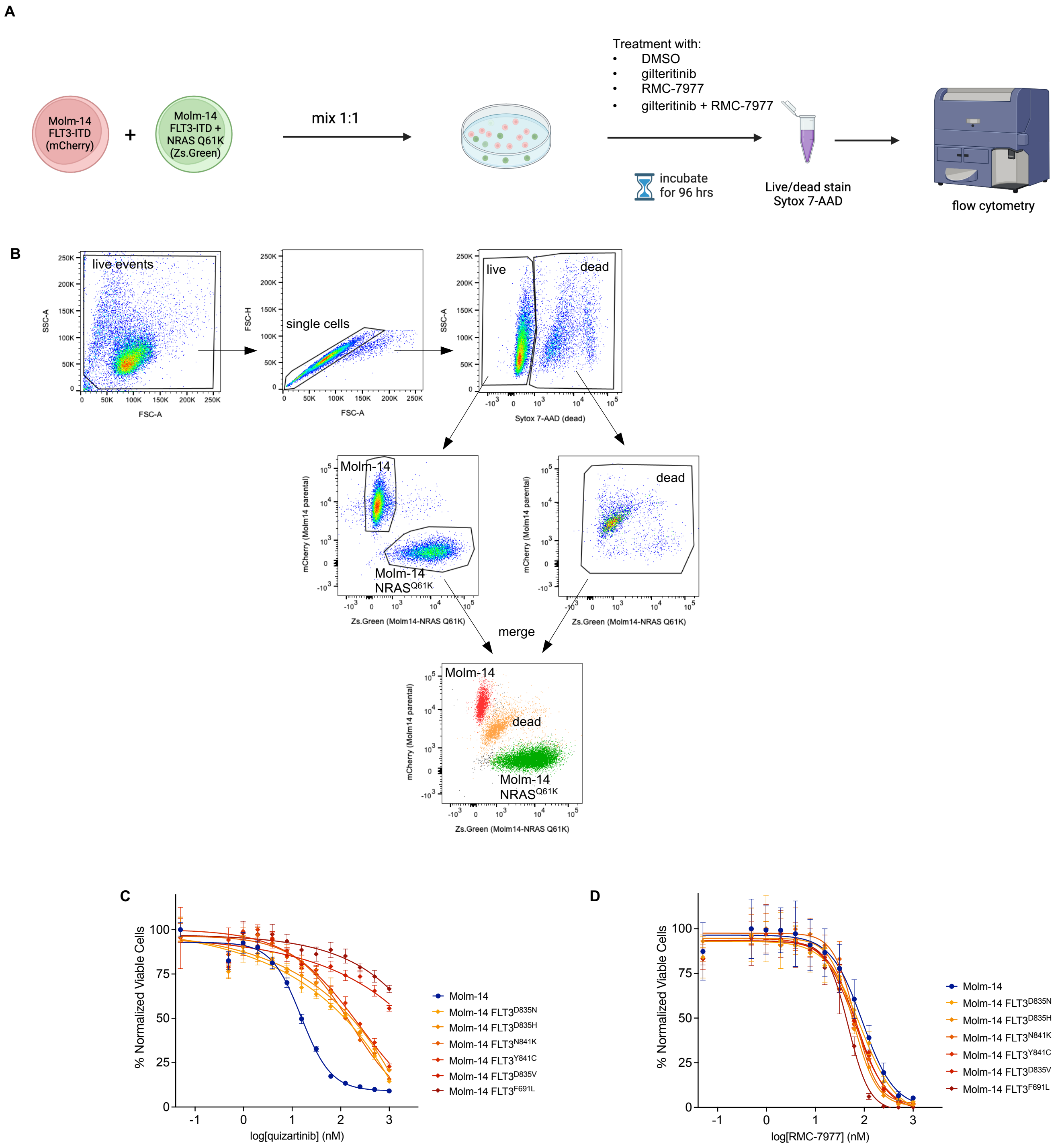
A
Molm-14
FLT3-ITD
(mCherry)
+
Molm-14
FLT3-ITD +
NRAS Q61K
(Zs.Green)
mix 1:1
Treatment with:
• DMSO
• gilteritinib
• RMC-7977
• gilteritinib + RMC-7977
incubate
for 96 hrs
Live/dead stain
Sytox 7-AAD
flow cytometry
B
live events
SSC-A
FSC-A
single cells
FSC-H
live
dead
Sytox 7-AAD (dead)
Molm-14
Molm-14
NRASQ61K
mCherry (Molm14 parental)
Zs.Green (Molm14-NRAS Q61K)
merge
Molm-14
dead
Molm-14
NRASQ61K
C
% Normalized Viable Cells
log[quizartinib] (nM)
Molm-14
Molm-14 FLT3D835N
Molm-14 FLT3D835H
Molm-14 FLT3N841K
Molm-14 FLT3Y841C
Molm-14 FLT3D835V
Molm-14 FLT3F691L
D
% Normalized Viable Cells
log[RMC-7977] (nM)
Molm-14
Molm-14 FLT3D835N
Molm-14 FLT3D835H
Molm-14 FLT3N841K
Molm-14 FLT3Y841C
Molm-14 FLT3D835V
Molm-14 FLT3F691L

**Supplementary Figure S3 – RMC-7977 overcomes resistance to FLT3 inhibitors.** A: Schematic representation of experimental workflow in Figure 3D. B: Gating strategy used for assay in Figure 3D. D-E: Concentration-response curves representing relative proliferation of Molm-14 parental cells or Molm-14 expressing secondary *FLT3* TKD mutations after 48 h of exposure to serial doses of either quizartinib (C) or RMC-7977 (D). Data represent the mean ± SD of three replicates.

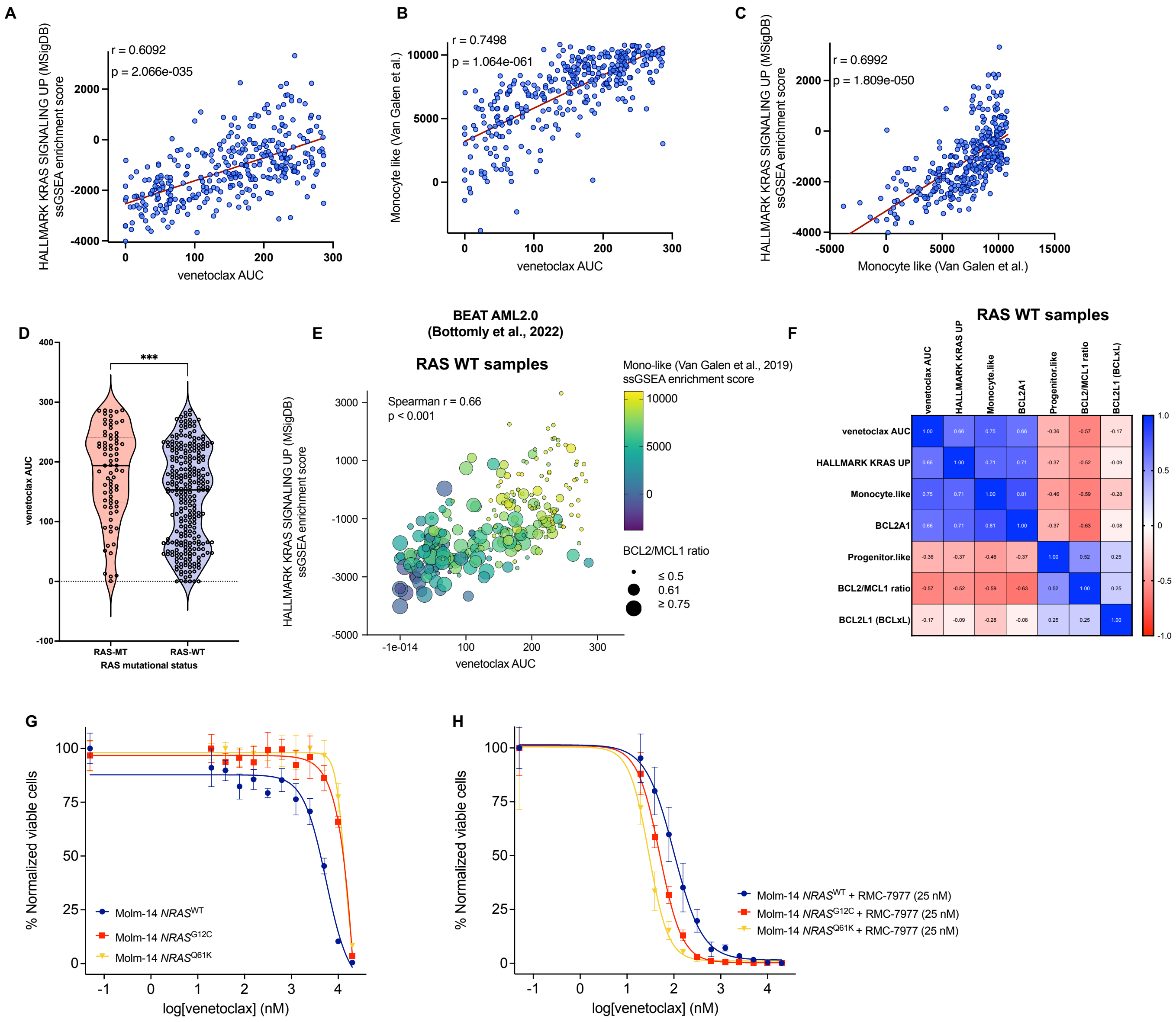
A
HALLMARK KRAS SIGNALING UP (MSigDB) ssGSEA enrichment score
r = 0.6092
p = 2.066e-035
venetoclax AUC
B
Monocyte like (Van Galen et al.)
r = 0.7498
p = 1.064e-061
venetoclax AUC
C
HALLMARK KRAS SIGNALING UP (MSigDB) ssGSEA enrichment score
r = 0.6992
p = 1.809e-050
Monocyte like (Van Galen et al.)
D
venetoclax AUC
***
RAS-MT
RAS-WT
RAS mutational status
E
BEAT AML2.0
(Bottomly et al., 2022)
RAS WT samples
Spearman r = 0.66
p < 0.001
HALLMARK KRAS SIGNALING UP (MSigDB) ssGSEA enrichment score
venetoclax AUC
Mono-like (Van Galen et al., 2019) ssGSEA enrichment score
BCL2/MCL1 ratio
≤ 0.5
0.61
≥ 0.75
F
RAS WT samples
venetoclax AUC
HALLMARK KRAS UP
Monocyte.like
BCL2A1
Progenitor.like
BCL2/MCL1 ratio
BCL2L1 (BCLxL)
G
% Normalized viable cells
Molm-14 NRASWT
Molm-14 NRASG12C
Molm-14 NRASQ61K
log[venetoclax] (nM)
H
% Normalized viable cells
Molm-14 NRASWT + RMC-7977 (25 nM)
Molm-14 NRASG12C + RMC-7977 (25 nM)
Molm-14 NRASQ61K + RMC-7977 (25 nM)
log[venetoclax] (nM)

**Supplementary Figure S4 – RMC-7977 targets RAS-mediated mechanisms of resistance to venetoclax in both $RAS^{WT}$ and $RAS^{mut}$ genetic backgrounds.** A-C: Correlations between *ex vivo* sensitivity to venetoclax (AUC), Hallmark *KRAS* ssGSEA enrichment score, and monocytic differentiation ssGSEA enrichment score based on BEAT AML2.0 transcriptomic and *ex vivo* drug sensitivity data. Nonparametric Spearman r coefficient and two-tailed p vales are shown. D: Difference in venetoclax sensitivity between $RAS^{WT}$ and $RAS^{mut}$ samples in BEAT AML2.0 trial. Unpaired two-tailed Mann-Whitney test was used for statistical significance (∗∗∗∗p ≤ 0.0001, ∗∗∗p ≤ 0.001, ∗∗p ≤ 0.01, ∗p ≤ 0.05, ns, non-significant). E: Bubble plot showing correlation between ex-vivo sensitivity to venetoclax (AUC) and RAS transcriptional activation (Hallmark KRAS Signaling Up, ssGSEA enrichment score) in 259 $RAS^{WT}$ primary AML samples in the BEAT AML trial. Size of bubbles represent BCL2/MCL1 normalized expression ratio and color gradient represents transcriptional scores of monocytic differentiation (Monocyte-like, ssGSEA enrichment score). F: Correlation matrix of nonparametric Spearman r correlation coefficients between venetoclax *ex vivo* sensitivity, ssGSEA enrichment scores of Hallmark *KRAS* Signaling Up, Monocyte-like, Progenitor-like, normalized expression of *BCL2A1* and *BCL2L1* and *BCL2*/*MCL1* expression ratio in $RAS^{WT}$ samples. G-H: Concentration-response curves representing relative proliferation of Molm-14 parental cells or Molm-14 expressing secondary $NRAS^{G12C}$ or $NRAS^{Q61K}$ mutations after 48 h of exposure to serial doses of venetoclax, in the absence (G) or presence (H) of RMC-7977 (25 nM).

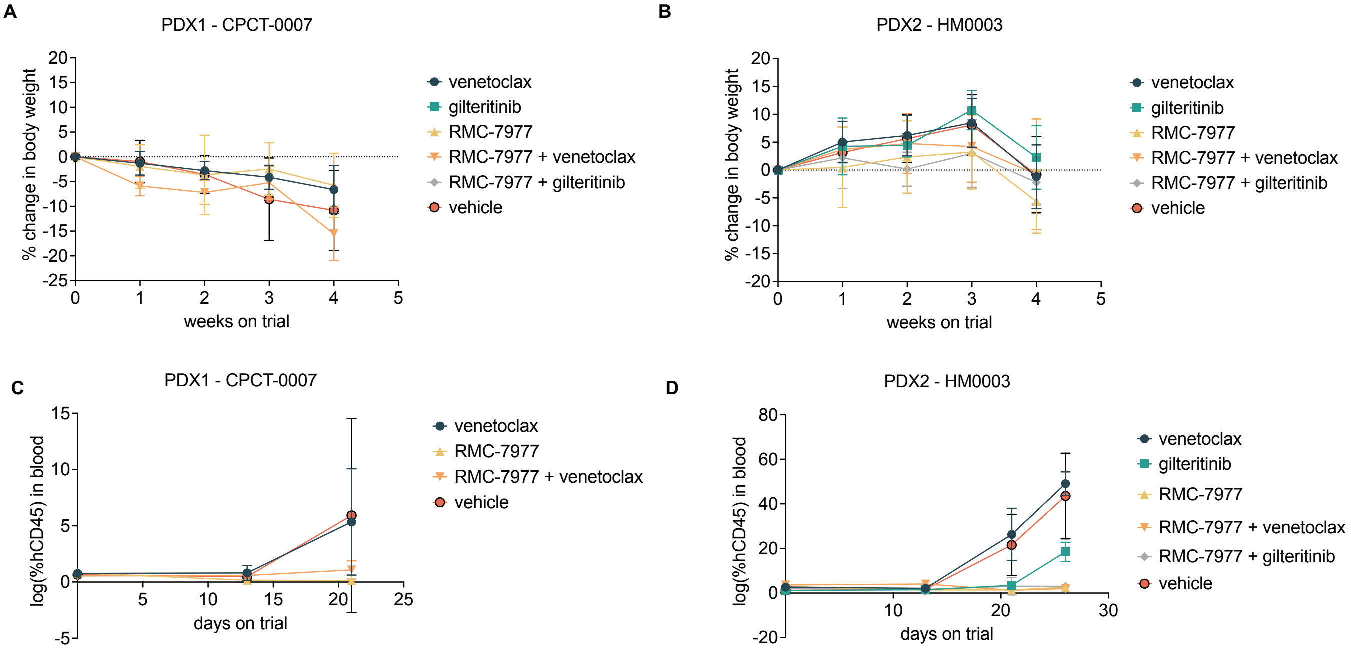

A
PDX1 - CPCT-0007
% change in body weight
weeks on trial
venetoclax
gilteritinib
RMC-7977
RMC-7977 + venetoclax
RMC-7977 + gilteritinib
vehicle
B
PDX2 - HM0003
% change in body weight
weeks on trial
venetoclax
gilteritinib
RMC-7977
RMC-7977 + venetoclax
RMC-7977 + gilteritinib
vehicle
C
PDX1 - CPCT-0007
log(%hCD45) in blood
days on trial
venetoclax
RMC-7977
RMC-7977 + venetoclax
vehicle
D
PDX2 - HM0003
log(%hCD45) in blood
days on trial
venetoclax
gilteritinib
RMC-7977
RMC-7977 + venetoclax
RMC-7977 + gilteritinib
vehicle

**Supplementary Figure S5 – In vivo activity of RMC-7977.** A-B: Body weight changes over the course of the PDX1 study (A) and PDX2 study (B). Data represent mean weight ± SD (n=6 mice/group). C-D: Quantification of hCD45+ cells over the course of treatment in the treatment groups in the PDX1 study (C) and PDX2 study (D).
